# Nutrient availability shapes the diversity and structure of microbial communities

**DOI:** 10.64898/2026.04.14.718562

**Authors:** Martina Dal Bello, Yizhou Liu, Akshit Goyal, Jeff Gore

**Affiliations:** Department of Ecology and Evolutionary Biology, Microbial Sciences Institute and Quantitative Biology Institute, Yale University, New Haven, CT 06511, USA; Physics of Living Systems Group, Department of Physics, Massachusetts Institute of Technology, Cambridge, MA 02139, USA; International Centre for Theoretical Sciences, Tata Institute of Fundamental Research, Bengaluru 560089, India

## Abstract

Nutrients are key drivers of microbial community structure, yet we lack a data-driven quantitative framework linking nutrient environments to community assembly. Here, using controlled microcosm experiments, we systematically probe the effects of nutrient number, concentration, and type on community diversity and structure. To explain these effects, we construct a minimal consumer-resource model incorporating resource competition and cross-feeding. We find that cross-feeding network structure is critical: only shallow, wide networks — where several byproducts are produced from a few supplied nutrients in a few trophic layers — reproduce a linear increase in diversity with the number of supplied nutrients. We also perform new experiments varying nutrient concentration and reveal that diversity slowly decreases with increasing concentration. We explain this finding by incorporating consumption-dependent toxicity into the model, consistent with spent-media measurements. This extended model not only recapitulates virtually all observed patterns but makes an independent prediction: at high nutrient concentrations, communities should be enriched in bistable species pairs, which we confirm experimentally. Our work demonstrates that minimal data-driven consumer-resource frameworks — systematically constrained by experiments — can unify and predict a broad range of nutrient–community relationships.

## Introduction

Microbial communities are ubiquitous across Earth’s biomes and play essential roles in ecosystem functioning: they constitute the basis of aquatic and soil food webs, mediate the decomposition of organic matter, and drive the biogeochemical cycling of carbon, nitrogen, phosphorus, and other elements [1]. In turn, the taxonomic and functional diversity of microbial communities is thought to underpin the stability and efficiency of these ecosystem processes. Understanding what controls microbial community diversity is therefore a central question in microbial ecology. Among the many environmental factors that shape microbial communities—including temperature [2, 3, 4], salinity [5, 6, 7], pH [8, 5, 9, 10], and moisture [11, 12]—nutrient availability stands out as a particularly fundamental driver, because it directly determines the metabolic niches accessible to community members and sets the energetic constraints under which species compete and help each other [13, 14, 15, 16, 17, 18]. Yet, despite decades of observational and theoretical work, we still lack a predictive understanding of how changes in nutrient availability shape the diversity and structure of microbial communities.

Nutrient pools in natural environments might differ along three main axes: their heterogeneity, i.e. the number of different compounds available for microbes to consume, their concentration, and type, for example whether available nutrients are sugars or organic acids, or labile vs. recalcitrant compounds. Surveys of natural microbiomes have revealed idiosyncratic responses of microbial community diversity to changes in nutrient heterogeneity and concentration. In temperate seas, winter mixing simultaneously replenishes surface nutrients and reshuffles deepwater taxa into the photic zone [19], so that nutrient-replete winter waters harbor more bacterial phylotypes than stratified, nutrient-poor summer waters in the Eastern Mediterranean [20] and Baltic Sea [7, 21, 22]. In contrast, increased nutrient availability close to roots compared to bulk soil decreases community diversity in the rhizosphere of plants over the course of microbial succession [14, 23]. Nevertheless, such studies struggle to separate the effect of nutrient availability from that of other environmental variables, preventing to causally connect observed patterns with their underlying processes.

Laboratory experiments, where nutrient pools can be finely tuned while other environmental variables are kept constant (Fig. 1A), are proving to be an ideal setting to probe the response of natural communities to changes in nutrient availability [24, 25, 13, 26, 27, 28, 29, 30]. Serial dilution of soil communities on single carbon sources yields assemblages whose family-level composition is reproducibly determined by nutrient identity [24, 26], with each additional supplied resource increasing species richness by only about one taxon [13]. In mixed-nutrient environments, a biomass-weighted sum of single-resource communities can provide a reasonable first approximation of community composition [31, 32], but systematic deviations emerge when the supplied resources are metabolically dissimilar, with one resource class dominating over the other [32]. In addition, it has been shown that assembly on complex substrates can be modular [28] or drive community divergence [29]. All these studies focused on nutrient identity and, while cross-feeding and nutrient-specific metabolic pathways can explain some of these results, a comprehensive predictive framework linking various axes of nutrient availability to community diversity is still lacking.

**Figure 1:**
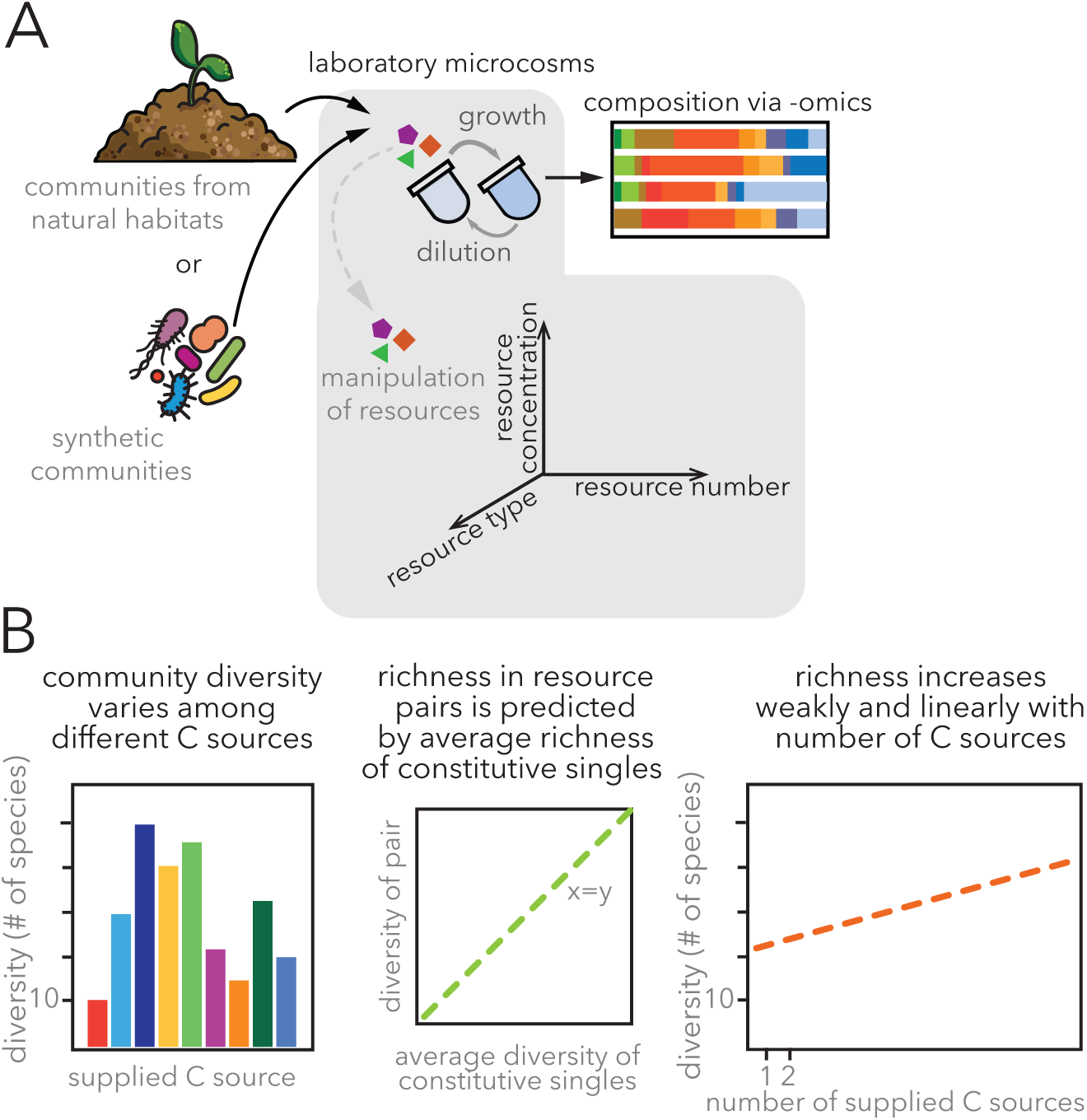
Community outcomes across across nutrient availability axes in laboratory microcosms. (A) Microbial communities from natural habitats (e.g., soil) or synthetic assemblages are cultured in laboratory microcosms to probe the effects of three axes of nutrient availability: resource type, resource number, and resource concentration. (B) Observed baseline diversity patterns derived from previous studies. Left: Community diversity varies significantly depending on the identity of the supplied single carbon source. Middle: The richness of communities grown on two resources is well-predicted by the average richness of communities grown on their constituent single resources. Right: Richness increases weakly and linearly with the number of supplied carbon sources.

Recently, consumer-resource models (CRMs) have emerged as powerful tools to understand the relationship between communities and their resource environments [33, 34, 35, 36, 37, 25, 38, 39, 40, 41]. These models typically incorporate ingredients such as resource competition, cross-feeding (the production of metabolic byproducts), and concentration-dependent growth rates. However, in virtually all these studies, modeling choices are *ad hoc* rather than systematic, and model ingredients are largely chosen to explain only a few (typically 1–2) observations of interest. As a result, models that can explain one set of patterns—say diversity—may fail to capture others—say those about community productivity [13, 25, 33, 24, 29]. Without a unified data-driven model that can simultaneously explain a large class of observed resourcecommunity relationships, our ability to predict and control community behaviors will remain severely limited.

Here we summarize patterns in bacterial community diversity observed in experiments controlling the number and type of carbon sources and uncover a new one by manipulating the concentration of individually supplied carbon sources. We then develop a consumer resource model highlighting the minimal ingredients needed to recapitulate the observed responses of community diversity to changes along the three nutrient availability axes.

## Results

### Community outcomes across nutrient availability axes in laboratory microcosms

Recent experimental studies manipulating the carbon sources supplied in the growth medium have revealed reproducible patterns in microbial community diversity. When different carbon sources, including sugars, organic acids and amino acids are supplemented as sole carbon sources in minimal medium, a range of community richness values are observed, both when natural communities and synthetic consortia are used as initial inoculum [24, 13, 42, 43, 25] (Fig. 1A). As the number of supplied carbon sources is experimentally increased, however, diversity increased slowly [13, 25], and in one instance [13] linearly with the number of resources. Intriguingly, in [13], the richness of communities grown on two resources was best predicted by the average richness of the communities grown on the constituent single resources. Finally, community evenness increased with the number of carbon sources [13, 25], showing that communities supported by a heterogeneous pool of resources exhibit a more balanced distribution of species’ relative abundances. As these studies noted, these patterns are quantitatively surprising. Existing models can neither capture all of them simultaneously, nor explain which ingredients are necessary for each pattern, e.g., for richness to increase linearly with the number of resources with a slope of roughly 1 [13].

Further, in these previous studies, the total concentration of the resources has been kept constant, leaving the response of communities to changes in resource concentration open. Hence, we set out to test the effect of varying the concentration of individually supplied resources on the diversity of bacterial communities. As in, we used laboratory microcosms inoculated with a microbial suspension with initially unknown diversity obtained from a soil sample. Each microbial microcosm is a well of a 96 deep-well plate filled with 300 *µ*L of minimal media supplemented with a single carbon source at four different concentrations, namely 0.001, 0.01, 0.1 and 1% w/v (Fig. 2A). We used a total of 12 carbon sources, including four sugars of different complexity, four organic acids, and four amino acids. Bacterial cultures were grown following a daily dilution protocol, whereby every 24 hours they were diluted 1/30X into fresh media. While several communities reached stability about four days after inoculation (Fig. S1), we passaged them for 7 days and measured the final community diversity as the number of amplicon sequence variants (ASVs) in each microcosm.

**Figure 2:**
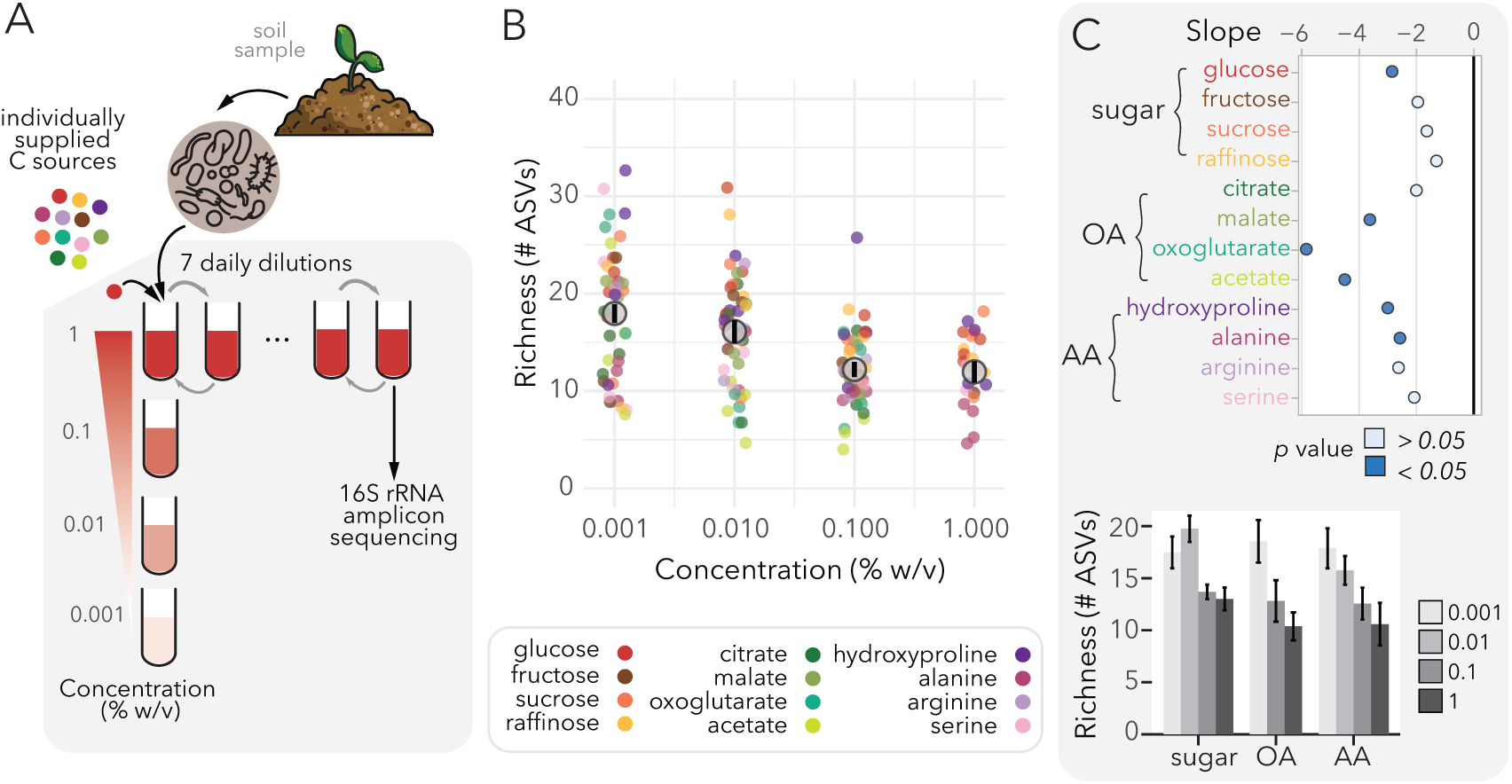
Increased carbon concentration drives a modest decrease in microbial community richness. (A) Experimental setup for concentration manipulation. Soil-derived communities were subjected to 7 daily dilution cycles in 12 individually supplied carbon sources at four concentrations (0.001%, 0.01%, 0.1%, and 1% w/v), followed by 16S rRNA amplicon sequencing. (B) Community richness (number of ASVs) across the four tested concentrations, showing a slight decrease as carbon concentration increases. (C) Top: Slope of richness versus concentration for individual sugars, organic acids (OA), and amino acids (AA). Data points are colored by statistical significance. Bottom: Average richness by resource class and concentration, highlighting a general decreasing trend, with sugars showing the largest difference between 0.01% and 0.1% w/v.

Confirming previous experimental studies [24, 13, 42], an average of 16.4 +/- 0.5 species coexisted in all individually supplied resources, regardless of their concentration. However, community richness slightly decreased as the concentration of the carbon sources increased in the growth media (Fig. 2B). This pattern was generally consistent across carbon sources, with 50% of them showing a statistically significant decrease in the number of species with the concentration of carbon (Fig. 2C). Sugars deviate slightly from this, showing the largest difference in richness between 0.01 and 0.1% w/v on average and no differences in richness between 0.1 and 1% w/v (Fig. 2D). Note that all communities grown in organic acids and arginine at 1% w/v were excluded (see Methods and Figs. S2, S6). A similar trend emerged when we looked at other diversity metrics (Figs. S7, S8) and was not driven by emergent co-limitation at the highest concentrations of the carbon sources (Fig. S9). These results are both surprising and counterintuitive: in standard consumer-resource models, increasing the supply concentration of a resource typically increases (and in some cases, does not change) the number of coexisting species [33, 13, 35, 24]. This is because a higher concentration of resource supply typically allows for more metabolic byproducts at higher concentrations, providing more energy to sustain additional species. The consistent decrease in richness we observe across carbon sources therefore suggests that high resource supply concentrations activate biological mechanisms not captured by current models — a point we return to below when developing our mathematical model.

### Shallow and wide cross-feeding networks reproduce linear increase in richness with nutrient number

Next, we sought to develop a consumer–resource model with the goal of finding the minimal set of “ingredients” needed to reproduce the observed experimental patterns. We started with a minimal model where species consume resources to grow and resources are supplied externally (see Methods for the full model development). Without cross-feeding, the competitive exclusion principle limits coexistence to at most one species per supplied resource, as confirmed by our simulations (Fig. 3B). Since a single resource can support dozens of species in our experiments, we introduced metabolic cross-feeding—the leakage of metabolic byproducts that become resources for other species [33, 24]—through the following set of ordinary differential equations:

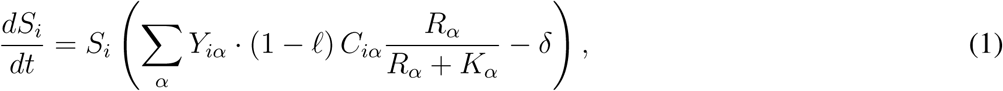

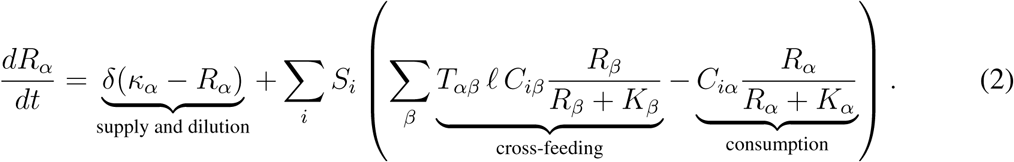

**Figure 3:**
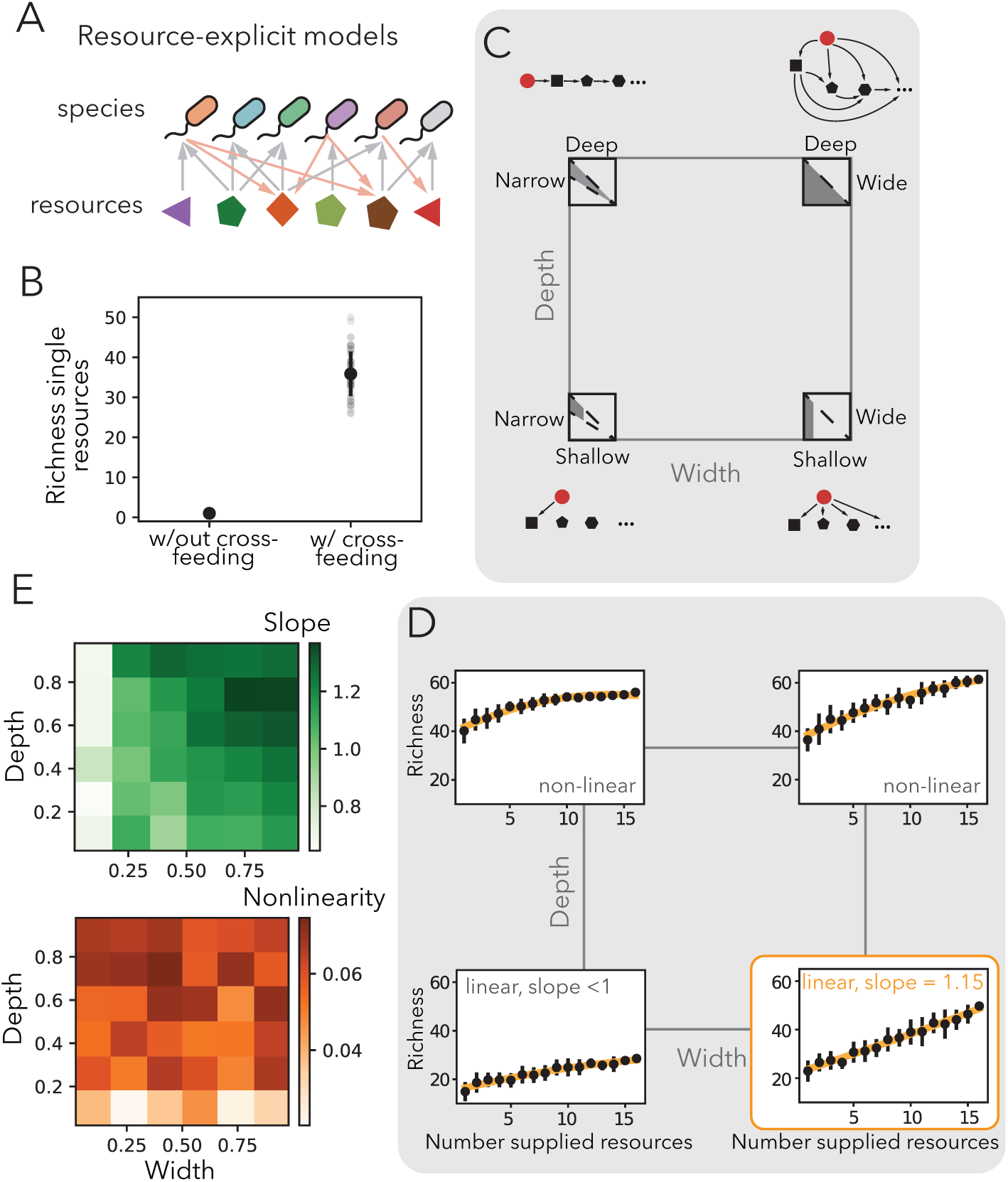
Shallow and wide cross-feeding networks are required to reproduce observed linear increase in richness with nutrient number. (A) Schematic of the minimal consumer-resource model incorporating species, resources, and metabolic cross-feeding. (B) Simulated richness on single resources demonstrating that cross-feeding is an essential ingredient for multi-species coexistence on a single supplied resource. (C) Systematic characterization of metabolic cross-feeding network structure across two axes: depth and width. (D) Heatmaps displaying the slope and nonlinearity of richness versus number of resources as a function of network width and depth. (E) Simulated community richness versus the number of supplied resources for the four extreme network architectures. Only shallow and wide networks successfully reproduce the experimentally observed linear increase in richness with a slope close to 1.

Here *S_i_* is the abundance of species *i*, *R_α_* is the concentration of resource *α*, *δ* is the dilution rate, *κ_α_* is the supply concentration, *C_iα_* is the consumption rate of species *i* on resource *α*, *K_α_*is the half-saturation constant, and *Y_iα_* is the yield. The parameter *ℓ* quantifies the fraction of consumption flux that is leaked as byproducts, and the matrix *T_αβ_* encodes the metabolic crossfeeding network, specifying how consumed resource *β* is converted to byproduct *α* (Fig. 3A). With cross-feeding, we could tune model parameters (see Methods) such that 20–30 species coexisted on a single supplied resource (Fig. 3B), consistent with our experimental observations.

While cross-feeding is sufficient to support high diversity on a single resource, not every network structure reproduces the observed linear scaling of richness with resource number [13]. To understand what constrains this relationship, we systematically varied the structure of the metabolic network along two axes: width, the number of distinct byproducts each nutrient generates; and depth, the number of trophic layers through which metabolites are further transformed (Fig. 3C). Deep networks produced a saturating, non-linear increase in richness, while narrow networks generated too few byproducts to substantially increase richness with more resources (Fig. 3D). Only shallow and wide networks—where each supplied nutrient generates several byproducts in few metabolic steps—reproduced the experimentally observed linear increase in richness with a slope roughly 1 (Fig. 3E, bottom right). To understand how network width and depth controlled this pattern, we fitted richness as a quadratic function of resource number for each network architecture and extracted the slope and nonlinearity (second-order coefficient; Methods). This revealed that network depth primarily controlled nonlinearity, while width set the slope (Fig. 3D). Intuitively, in shallow and wide networks each newly supplied resource generates several non-overlapping byproducts, consistently opening new niches for additional species without saturating the available metabolic niche space. A shallow and wide cross-feeding architecture is therefore both necessary and sufficient to quantitatively reproduce the observed linear increase in richness with the number of supplied resources.

### Wide, shallow cross-feeding networks capture nutrient number patterns but are insufficient for concentration patterns

To evaluate the predictive power of the cross-feeding network we identified, we tested whether it could help capture other key experimental relationships regarding the number of available nutrients. Remarkably, without any further parameter tuning, the model successfully recapitulated the remaining three observed trends. First, the simulated richness of communities grown on pairs of resources was roughly the average richness of the constituent single-resource communities (Fig. 4B), mirroring our experimental observations. Second, the model correctly predicted that the total community biomass saturates stays roughly constant with the number of supplied resources (Fig. 4C). Third, the model also reproduced the observed increase in community evenness, measured using rank-abundance curves (Fig. 4D).

**Figure 4:**
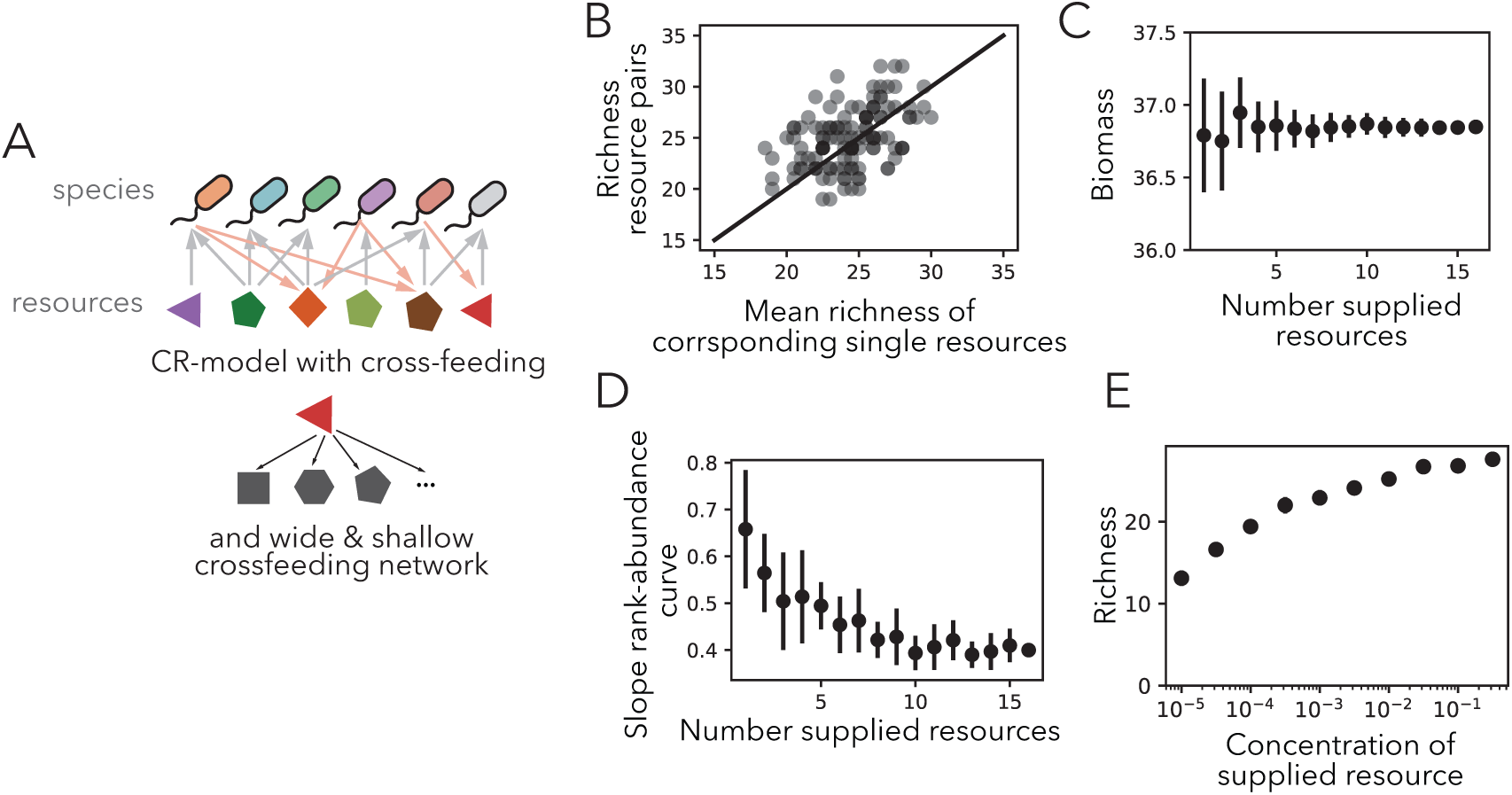
Model with wide and shallow cross-feeding networks captures nutrient number patterns but fails at high concentrations. (A) Schematic of the standard consumer-resource model featuring the identified wide and shallow cross-feeding network. (B) Simulated richness of resource pairs strongly roughly matches with the mean richness of corresponding single resources. (C) Simulated community biomass as a function of the number of supplied resources. (D) Slope of the rank-abundance curve versus the number of supplied resources. (E) Simulated richness increases with the concentration of the supplied resource. This directly contradicts the experimental findings in Fig. 2, revealing a missing ingredient in the model.

Despite these successes in capturing the effects of nutrient type and number, the model failed to predict changes in community diversity with increasing nutrient concentration. Specifically, the model predicted an increase in community richness with resource concentration (Fig. 4E), in contrast with our experimental results where richness decreases with concentration (Fig. 2B). This qualitative discrepancy reveals that our current model is missing something important at high nutrient supply concentrations which hinders coexistence.

### Spent media of species grown at highest concentration of carbon is harmful

To understand the effect of increasing supply concentration on growth in a community, we performed experiments where we grew isolates on their spent media. Specifically, we isolated about 40 species from equilibrated communities in the experimental microcosms, and chose 8 that were prevalent in glucose, spanned more than one phylum, and could be distinguished based on their colony morphology. Specifically, we picked six representatives of the phylum Proteobacteria (*Raoultella terrigena*, *Klebsiella*, *Agrobacterium*, *Pseudomonas*), one Bacteroidota (*Sphigobacterium*), and one Actinobacteriota (*Paenarthrobacter*) (Fig. 5A). We grew each species in minimal medium supplemented with glucose at the four concentrations included in the community experiment for 24 hours, collected the spent media and re-supplemented it with the original glucose concentration and components of the minimal medium. We then grew all the 8 species for another 24 hours (Fig. 5B). At the end of the 24 hours, we measured the optical density (OD) of each culture to assess the impact of secreted metabolites at different glucose concentrations on the growth of the 8 community members. Since overall growth increased with glucose concentration, we rescaled the OD for species at each concentration to allow for a clearer visual comparison (Methods). We found that spent media of species grown at 0.001, 0.01 and 0.1 % w/v of glucose did not harm nor facilitate the growth of community members (Fig. 5C). By contrast, the spent media collected after growth at glucose 1% w/v harmed most species (Fig. 5C), with more than 30% of species experiencing a reduction in final optical density of more than 20% (Fig. S10). Thus, our experiments show that isolates grow to much lower abundances at the highest supply concentrations. This suggests that at such high concentrations, there is a shift in the secreted metabolites towards toxic compounds.

**Figure 5:**
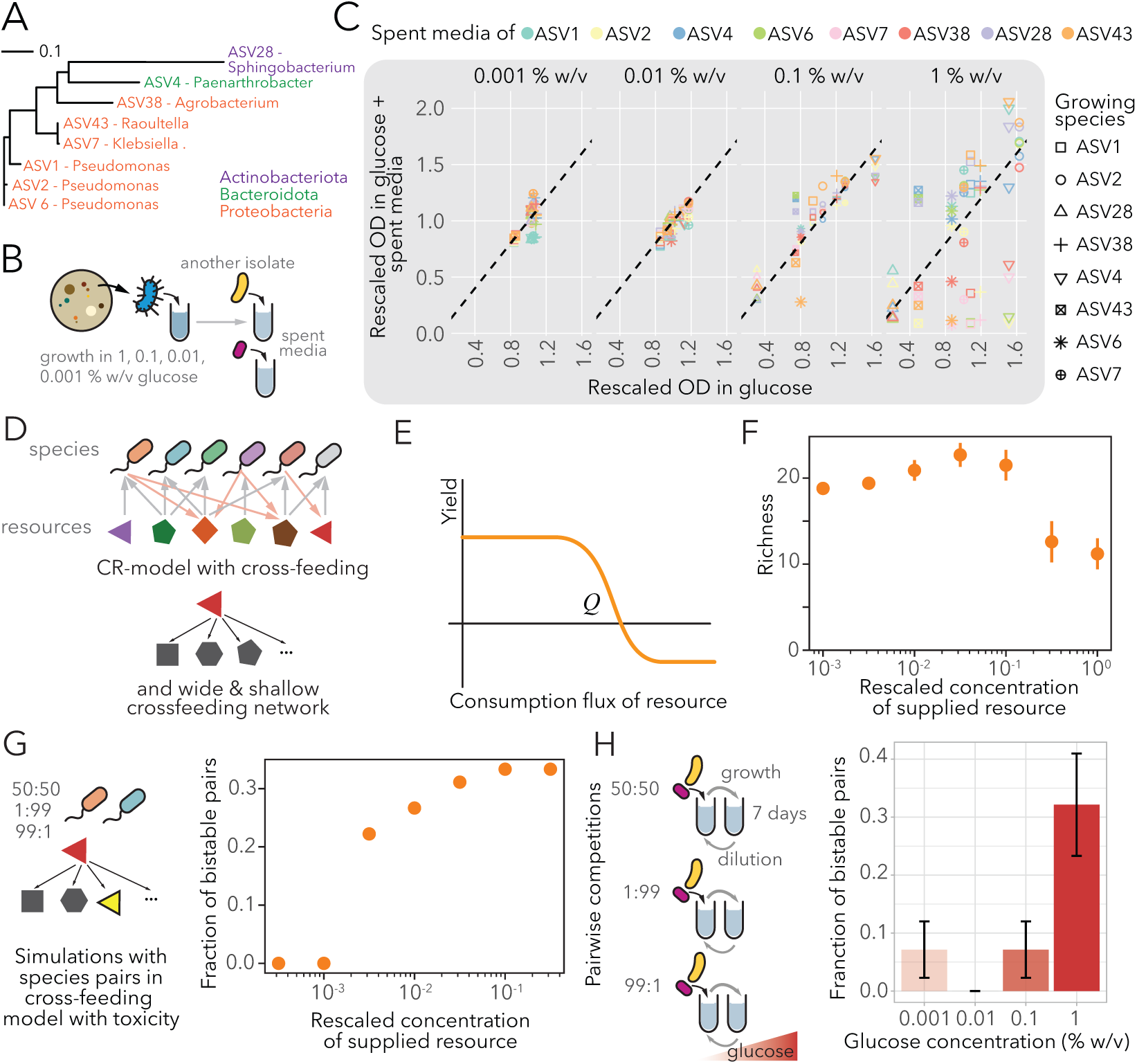
Consumption-dependent toxicity, suggested by spent-media experiments, explains diversity decline at high nutrient concentrations. (A) Phylogenetic tree of eight representative isolates spanning multiple phyla, selected from the experimental microcosms for spent-media assays. (B) Experimental design for the spent-media assay. Isolates were grown in spent media collected from cultures previously grown at varying glucose concentrations. (C) Growth (rescaled OD) of isolates in fresh glucose versus glucose supplemented with spent media. Spent media from the 1% w/v glucose condition significantly hindered growth for most species, indicating the accumulation of toxic byproducts. (D) Updated consumer-resource model schematic incorporating concentration-dependent toxicity. (E) Model function where species yield transitions from positive (growth-promoting) to negative (toxic) once the consumption flux exceeds a critical threshold *Q*. (F) Simulated richness versus rescaled concentration under the updated model, successfully recapitulating the experimentally observed decrease in diversity at high nutrient concentrations. (G) Independent prediction: simulating pairwise competitions using the updated model at increasing concentrations revealed a sudden jump in the fraction of pairs with bistable outcomes, (H) confirmed by pairwise competition experiments. The binomial standard error is displayed (*n* = 28 for each glucose concentration).

### Consumption-dependent toxicity reproduces diversity decrease with nutrient concentration

Inspired by these experiments, we sought to incorporate consumption-dependent toxicity into our model (Fig. 5D). To do this in a minimal way, we assumed that species growth yields *Y* ^+^ were now a function of the consumption flux (in our earlier model, they were fixed). Specifically, as the consumption flux of a nutrient increased beyond a threshold flux *Q*, yields switched from positive (promoting growth) to negative (harming growth; Fig. 5E; see Methods for details). This mimicked the spent-media experiments where the reduction in growth occurred only at the highest supply concentrations (Fig. 5F). For simplicity, we assumed the same consumption flux-dependence for all species (we later relax this assumption in the SI; see Fig. S11).

This simple and minimal addition to the model successfully reconciled predictions with our experimental observations. The refined model now predicted a decrease in community richness at high supply concentrations (Fig. 5C), as observed in our experiments (Fig. 2B). This decrease in richness can be understood by noting that increasing nutrient supply increases consumption across all species, eventually triggering toxicity which kills the most sensitive species. Crucially, because toxicity in our model only “kicks in” at high concentrations, the modified model can still successfully predict all other patterns regarding nutrient number and type (seen in Figs. 3 and 4; also see Fig S12). Thus, by integrating resource competition with structured cross-feeding and consumption-dependent toxicity, we now have a model that can quantitatively recapitulate the observed nutrient-community relationships across all three axes: nutrient number, type and concentration.

### Combined model predicts that communities at high nutrient concentrations should be enriched in bistable species pairs

A key test of any data-driven model is whether it can make predictions beyond the data used to constrain it. Analysis of our model at high supply concentrations (see Methods for details) revealed a striking prediction: the fraction of species pairs exhibiting bistability—where the surviving species depends on the initial inoculum—increased dramatically at the highest nutrient supply concentrations (Fig. 5G). This occurs because consumption-dependent toxicity introduces strong effective inhibition between species sharing similar metabolic niches, so that the initially more abundant species pollutes the environment and significantly affects the growth of the other.

To test this prediction, we used the 8 isolates previously selected for the spent media assays and performed all 28 pairwise coculture experiments at varying glucose concentrations. To test for emergence of bistability at steady state, cocultures were started at three different initial fractions (the two species inoculated in equal abundance, at an abundance rate of 99 to 1%, and 1 to 99%) and passaged them for 7 days following the same protocol used for the community assembly experiments. Consistent with the model’s prediction, we found that roughly 30% of species pairs grown at 1% w/v carbon were bistable—where the winner depended on initial composition—compared with almost no bistable pairs when grown at lower concentrations (Fig. 5H).

## Discussion

Over the past several years, high-throughput enrichment cultures, where natural microbial communities are serially passaged in defined synthetic media under controlled conditions, have emerged as a powerful experimental platform for studying microbial community assembly [27]. Analogous approaches where strains are mixed to obtain synthetic communities provide control also over taxon identity and number but at the expenses of realism [25, 42]. A striking feature of this body of work is the consistency of its findings across independent laboratories and inocula. A range of carbon sources when supplied as the sole resource sustain stable multispecies communities, defying the principle of competitive exclusion [24, 13, 26, 42, 25, 43]. When resources are combined in mixtures of increasing complexity, diversity increases slowly [13, 25]. These patterns have been observed with different soil inocula or synthetic communities, different carbon source panels, and in different laboratories, pointing to general ecological principles at work. Yet, despite this consistency, we lacked a unified quantitative framework that could simultaneously explain all observed patterns. Here, building on published results and new results about the response of community diversity to changes in nutrient concentration, we provide such synthesis by developing a consumer resource model model that is iteratively constrained by the data across all three nutrient axes (number, type, and concentration). Our model not only recapitulates the full set of observed patterns but generates an independent prediction, the nutrient concentration -dependent bistability, that we confirm experimentally. This progression from consistent empirical patterns to a minimal, data-driven predictive model illustrates the power of enrichment cultures as a platform for building quantitative theory in microbial ecology.

Consumer-resource models have become central tools in microbial ecology [33, 34, 35, 36, 37, 25, 38, 39, 40, 29, 41, 44], yet most studies validate against 1–2 empirical patterns at a time. Here, we used the richness–resource-number relationship to discriminate among entire classes of cross-feeding network architectures, performing model selection directly constrained by data. The shallow and wide architecture we identify, where nutrients are converted into several distinct byproducts in few biochemical steps, is independently supported by exometabolomics studies showing that bacteria simultaneously secrete multiple central metabolic intermediates [45], and by genome-scale modeling predicting that costless metabolite secretion stabilizes after a single metabolic step [46]. Biologically, this is also consistent with overflow and mixed-acid fermentation, where rapid growth generates diverse partial oxidation products [47]. We also previously used a similar cross-feeding network architecture, which we discovered using a scope expansion analysis on a “universal” network of microbial metabolism [13]. Our work thus suggests that macroscopic patterns, such as richness scaling with resource number, can be used to infer mesoscopic properties of the community metabolic network.

Our results reveal a sharp contrast between two axes of nutrient availability: increasing the number of carbon sources expands community diversity, while increasing the concentration of any single resource modestly reduces it. This parallels the ecological distinction between niche dimensionality and resource enrichment, but here we provide mechanistic explanations grounded in a data-constrained model. The linear richness–resource number relationship arises from cross-feeding: each additional carbon source generates new metabolic byproducts, expanding available niches roughly proportionally. This contrasts with plant communities, where adding a limiting nutrient reduces diversity [48]. In microbial communities, the relevant comparison is increasing the supply of an existing resource—which, as we show, shifts metabolism towards toxic byproducts and reduces diversity. These results suggest that the chemical diversity of the resource pool, rather than its energy content, is a primary driver of microbial coexistence. The concentration-dependent diversity loss we observe parallels the classical paradox of enrichment [49] and is consistent with recent experiments showing that high nutrient concentrations promote negative ecological interactions among bacteria [50, 51]. In those studies, however, toxicity derived primarily from strong changes in the pH of the growth medium [50, 51] and could be removed by buffering the environment [52, 9]. Our spent-media experiments suggest a more general mechanism: only in one instance—*Klebsiella sp.* at the highest glucose concentration—could we trace growth inhibition to environmental acidification (Fig. S13). In the remaining cases the toxic compounds remain unidentified, though plausible candidates include secondary metabolites [53, 54], redox-active metabolites [55, 56], and overflow products that accumulate to inhibitory concentrations [47, 57]. In our model, toxicity emerges when the consumption flux exceeds a critical threshold, requiring only that certain byproducts—benign at low concentrations—become harmful as they accumulate [47, 57]. The same phenomenology would arise if entirely new toxic compounds were produced at high concentrations, such as rhamnolipid biosurfactants in *Pseudomonas aeruginosa* [58]. Our spent media experiments cannot distinguish between these two scenarios, and targeted metabolomics across concentrations will be needed to resolve this question. Interestingly, some species in our spent-media experiments actually grew better due to spent media at the highest glucose concentration (Fig. 5C). This is naturally explained by our model: because consumption fluxes vary across species, some species with lower consumption fluxes may remain below the toxicity threshold and actually benefit from the cross-fed byproducts present in the spent medium.

Finally, our model predicted that increasing nutrient concentration results in more species pairs exhibiting bistability, which our experiments confirmed. Our model also predicts multistability in the full communities (Fig. S14). Our enrichment experiments did not have enough replicates to test this conclusively. Nevertheless, communities grown on glucose at the highest concentration did exhibit two different biomass states — a high-biomass state when *Pseudomonales* dominate and a low-biomass state where *Klebsiella* dominates (Fig. S15). Both in communities and pairs, toxicity generated at high nutrient concentrations might be frequency-dependent: whichever species has an initial advantage pollutes the shared environment harming the other competitors that share a similar niche. This creates the positive feedback loops that generate bistability, a mechanism analogous to the inhibition-driven priority effects described in *Streptomyces* communities [59] and to pH-mediated environmental modification in two-species communities [9, 60]. Our experimental results that roughly 30% of species pairs become bistable at the highest nutrient concentration connects microbial metabolism to a broader ecological pattern: nutrient enrichment has long been recognized as a driver of catastrophic regime shifts in lakes, grasslands, and coral reefs [61]. More broadly, our results show that the three axes of nutrient availability (number, type, and concentration) shape microbial communities through distinct but interconnected mechanisms: niche creation via cross-feeding, niche identity via metabolic specificity, and niche collapse mediated by toxicity. By identifying these mechanisms within a single quantitative framework, our work provides a foundation for predicting how changes in nutrient environments will reshape microbial communities across both laboratory and natural ecosystems.

## Acknowledgements

We would like to thank Shaul Pollak and Jacopo Grilli for helpful discussions, and Abheepsa Nanda and Jana Huisman for help with isolating and sequencing bacterial isolates. AG acknowledges support from the Ashok and Gita Vaish Junior Researcher Award, the ANRF Ramanujan Fellowship, as well the DAE, Govt. of India, under project no. RTI4001.

## Methods

### Growth media preparation

All chemicals were purchased from Sigma-Aldrich unless otherwise stated.

All bacterial cultures were grown in M9 minimal media (prepared from 5X M9 salts, 1000X Trace Metal Mixture (Teknova) and 1M stock solutions of MgSO_4_ and CaCl_2_) supplemented with one of twelve carbon sources (glucose, fructose, sucrose, raffinose, citric acid, malate, acetate, trans-4-hydroxy-d-proline, arginine, alanine, serine) at four different concentrations (0.001, 0.01, 0.1 and 1% w/v). To obtain the different concentrations 10X stock solutions of each carbon source was prepared prior to the experiment by dissolving 100 gr/L of each carbon source in 500 ml of ddH2O and diluting 10-fold to obtain stocks at 10, 1, 0.1 and 0.01 % w/v. Each stock solution was filter-sterilized with a 0.22 *µ*m filter and kept on the bench. M9 minimal medium was prepared fresh daily.

### Collection of microbial communities from the environment

The soil from which the initial inoculum was obtained was sampled from a lawn in Cambridge, Massachusetts, at a depth of about 15cm using a sterile corer and tweezers. Once in the lab, a total of 2g of the collected soil was diluted in 40ml phosphate-buffered saline (PBS, Corning), then vortexed at intermediate speed for 30s and incubated on a platform shaker (Innova 2000, Eppendorf) at 250 r.p.m. at room temperature. After 1 h, the sample was allowed to settle for ∼5min and the supernatant was filtered with a 100*µ*m cell strainer (Thermo Fisher Scientific) and then directly used for inoculation. Both the original soil sample and the remaining supernatant were stored at 80°C for subsequent DNA extraction.

### Experimental microcosms

Aliquots (10*µ*L) of the supernatant containing the soil microbial suspension were inoculated into 290*µ*L growth media in 96-deepwell plates (deepwell plate 96/500*µ*L, Eppendorf), for a total of 206 microcosms (4 replicates for each resource and concentration in addition to 7 water and 7 M9 without carbon inocula). Deepwell plates were covered with AeraSeal adhesive sealing films (Excel Scientific). Bacterial cultures were grown at 30°C under constant shaking at 1,350 r.p.m. (on Titramax shakers, Heidolph). To avoid evaporation, they were incubated inside custom-built acrylic boxes.

Every 24 h, the cultures were thoroughly mixed by pipetting up and down three times using the VIAFLO 96-well pipettor (Viaflo 96, Integra Biosciences; settings: pipette/mix program aspirating 10*µ*L, mixing volume 100*µ*L, speed 6) and then diluted 1/30× into fresh media. We applied a total of 7 daily dilution cycles. At the end of every cultivation day, we measured the optical density (OD600) using a microplate reader (Tecan Nano). The remaining bacterial culture was frozen at 80°C for subsequent DNA extraction.

### DNA extraction, 16S rRNA sequencing and analysis pipeline

DNA extraction was performed with the Zymo Quick-DNA Fungal/Bacterial 96 Kit following the provided protocol. The obtained DNA was used for 16S amplicon sequencing of the V4 region. Library preparation and sequencing, which was done on an Illumina MiSeq platform, were performed by the Massachusetts Institute of Technology BioMicroCenter.

We used the R package DADA2 [62] to obtain the amplicon sequence variants (ASVs) following the workflow described in Callahan et al [63]. Taxonomic identities were assigned to ASVs using the SILVA version v138.1 database. Relative abundances were corrected by 16S rRNA copy number using rrnDB database version 5.8 [64].

### Soil strain isolation, sequencing, and identification

On day 7, cultures were plates on 20% LB (Beckton Dickinson) agar (bacteriological agar VWR) plates (150×15mm) and left to grow at 30 °C for two days. Representative colonies displaying different morphologies were picked and individually streaked onto 100×15mm LB plates. Single colonies were then picked from these plates and grown in 5 ml LB broth overnight. 1 ml of culture was then combined with 1 ml sterile 60% glycerol solution and stored at −80°C.

Isolates were streaked onto fresh 20% LB agar plates from the −80 °C stock and sent for 16S rRNA gene Sanger sequencing with Azenta Life Sciences. We used their in-house pipeline to trim and merge the forward and reverse reads. We assigned taxonomy using DADA2 and the SILVA database (v138.1). We aligned the 16S genes with MAFFT [65], and used RaxML [66] with a GTR model with Γ site variation to estimate a maximum likelihood phylogeny for all isolates. ASVs were matched to isolate 16S rRNA gene sequences using BLAST at *>* 97% identity [67].

### Spent media experiments

Among the isolated strains, we picked 8 that were abundant in glucose and spanned more than one phylum. Six representatives of the phylum Proteobacteria (Raoultella terrigena, Klebsiella, Agrobacterium, Pseudomonas), one Bacteroidota (Sphigobacterium), and one Actinobacteriota (Paenarthrobacter) were used in spend media experiments. To generate spent media, bacterial cultures were first grown for 24 hours at 30°C in 5 ml LB medium from a single colony with shaking at 250 rpm on a platform shaker. Cultures were then washed twice with 5ml PBS. Subsequently, 5 *µ*L of each washed LB culture was used to inoculate 5 ml of M9 minimal medium supplemented with glucose at four concentrations (1%, 0.1%, 0.01%, and 0.001% w/v) in 50 ml falcon tubes, prepared by combining M9 minimal medium with 10X glucose stocks to achieve the final concentrations. These 32 cultures were incubated for 24 hours at 30°C with shaking at 250 rpm. Spent media were obtained by centrifuging the glucose cultures at 3,220 × g for 10 minutes, collecting the supernatants, and filter-sterilizing them through 0.22 *µ*m membranes (Steriflip Filtration Unit, MilliporeSigma). Sterility was verified by spotting 50 *µ*L of each spent medium onto LB plates. Spent media were supplemented with glucose at its original concentration before dispensing them into 96-well plates and inoculating them with 3*µ*L of washed and 1000-fold diluted LB culture of each isolate (started 24h before), yielding a total of 256 spent media cultures. For comparison, each species was grown in fresh M9 supplemented with glucose at four concentrations (1%, 0.1%, 0.01%, and 0.001% w/v) in the same plates. Optical density (OD) was measured after 24 hours in a microplate reader (BMG Spectrostar Nano).

For visualization purposes, optical densities for each species, spent media, and glucose concentration were rescaled by dividing each value by the OD observed in the corresponding glucose concentration.

### Pairwise competitions

We used the 8 isolates described above for the spent media assays to set up pairwise competitions at four concentrations of glucose. Each strain was initially grown for 24 hours at 30°C in 5 ml LB medium from a single colony with shaking at 250 rpm on a platform shaker. Cultures were than wash twice in 5ml of PBS, diluted to the OD corresponding to a starting density of 5*106^^^ CFU/mL (based on on OD/CFU mapping), and subsequently mixed to obtain the intended starting ratios (50:50, 99:1, 1:99). 10 *µ*L of each strain-pair mixture were inoculated into 96 well 500 *µ*L deepwell plates (Eppendorf) with 290 *µ*L M9 supplemented with either 1%, 0.1%, 0.01% or 0.001% w/v glucose per well (for a total of 4 plates, one for each concentration). The four plates were incubated at 30°C on a benchtop shaker (Titrama× 100) at 1350 rpm. For seven consecutive days, we diluted the culture in each well 1:30 and transferred the communities to a plate with fresh culture medium (using the Integra VIAFLO96). After every transfer, we used 100 *µ*L from the old plate to measure OD600 (Tecan Infinite M Nano). At days 1, 3, 5 and 7, we diluted each plate in PBS to 10^−6^ and 10^−7^ (for glucose 1%), 10^−5^ and 10^−6^ (for glucose 0.1%), 10^−4^ and 10^−5^ (for glucose 0.01%), 10^−3^ and 10^−4^ (for glucose 0.001%), spot-plated 10 *µ*L per well, and counted the corresponding colonies after 2-3 days.

Pairs were considered bistable when we observed different stable outcomes depending on the initial starting ratio (50:50, 99:1, 1:99).

### Data analysis

Unless otherwise stated, analyses were conducted in R version 4.5.3. Sequencing data was handled using the R package phyloseq [68]. Community diversity was also measured using the Shannon diversity index and the Shannon entropy index following refs [69, 70].

### Mathematical modeling

#### Consumer-resource model

We modeled microbial community dynamics using a consumer– resource framework in which species grow by consuming resources that are supplied externally in a chemostat-like setting. The dynamics of species abundance *S_i_* and resource concentration *R_α_*are governed by

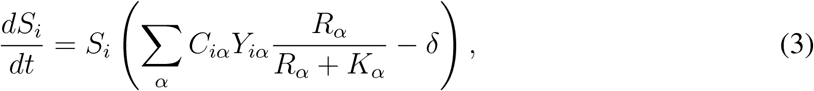

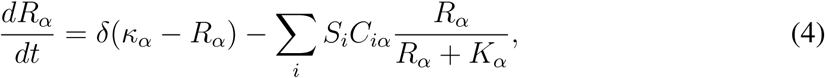

where *δ* is the dilution rate, *κ_α_* is the supply concentration of resource *α*, *C_iα_* is the consumption rate of species *i* on resource *α*, *K_α_* is the half-saturation constant, and *Y_iα_*is the yield relating resource consumption to biomass production. Species were allowed to consume multiple resources.

#### Cross-feeding

To allow more species to coexist than the number of supplied resources, we introduced metabolic cross-feeding following Ref. [33], respecting mass conservation:

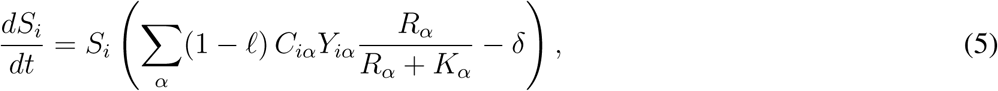

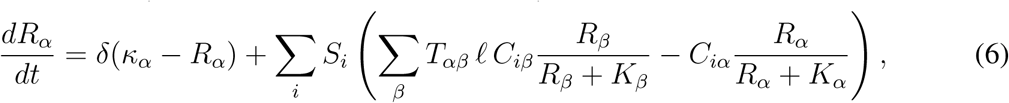

where *ℓ* is the fraction of consumed resource mass leaked as metabolic byproducts and *T_αβ_* specifies the fraction of consumed resource *β* converted to resource *α*, subject to the conservation constraint Σ*_α_T_αβ_* = 1.

#### Metabolic network structure

We restricted the metabolic network to be loop-free by imposing *T_αβ_* = 0 for *α* ≤ *β*, so that resources with smaller indices correspond to nutrients and those with larger indices to waste products. We parameterized the network along two structural axes. The *width* controls how many distinct byproducts each nutrient can yield: we set *T_αβ_*= 0 for *α* − *β* exceeding a threshold equal to *β* multiplied by a width fraction. The *depth* controls how many resources can serve as substrates for further metabolism: we set *T_αβ_* = 0 for *β* larger than a depth fraction multiplied by the total number of resources *M*. Non-zero entries of *T_αβ_* were drawn uniformly at random and renormalized to satisfy the conservation constraint.

#### Consumption-dependent toxicity

To capture the experimentally observed decrease in diversity at high nutrient concentrations, we modified the yield *Y_iα_* to depend on the consumption flux rather than a constant. Specifically, the yield switched from positive (growth-promoting) to negative (growth-inhibiting) when the consumption flux *C_iα_R_α_/*(*R_α_* + *K_α_*) exceeded a threshold *Q*. For simplicity, the same threshold was used for all species and resources. Mathematically,

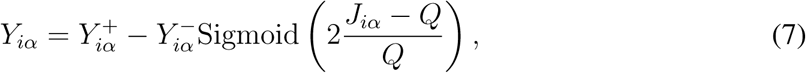

where 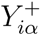 and 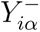 are constant. Based on the design, we have a transition of *Y_iα_* happening roughly in the range from *Q/*2 to 3*Q/*2.

#### Simulations

All simulations were performed with a species pool of *N* = 100 and a resource pool of *M* = 200. Consumption rates *C_iα_*were drawn from a truncated Gaussian distribution, where we first sample a Gaussian variable with mean zero and standard deviation 2, and then keep it if positive and set it to zero if the variable is negative. To assess richness, we numerically integrated the dynamical equations to a steady state and counted the number of species with abundance above a threshold of 10^−5^. The dilution rate *δ* and leakage *ℓ* were scanned at first and kept as *δ* = 0.1 and *ℓ* = 0.8, such that with cross-feeding, a single resource can support around twenty species, similar to experiments. When varying the number of supplied resources, we sequentially set *κ_α_* ≠ 0 starting from the smallest resource indices. When varying the concentration, we scaled *κ_α_* over four orders of magnitude. After introducing toxicity, we also scanned 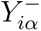 and *Q* systematically and chose the combination such that richness decreases in a range of concentration with a similar span in log scale as the empirical results. For bistability analysis, we simulated all species pairs at varying supply concentrations with two different initial conditions (each species initially dominant) and classified a pair as bistable if the two simulations converged to distinct steady states.

## Supplementary Figures

**Figure S1:**
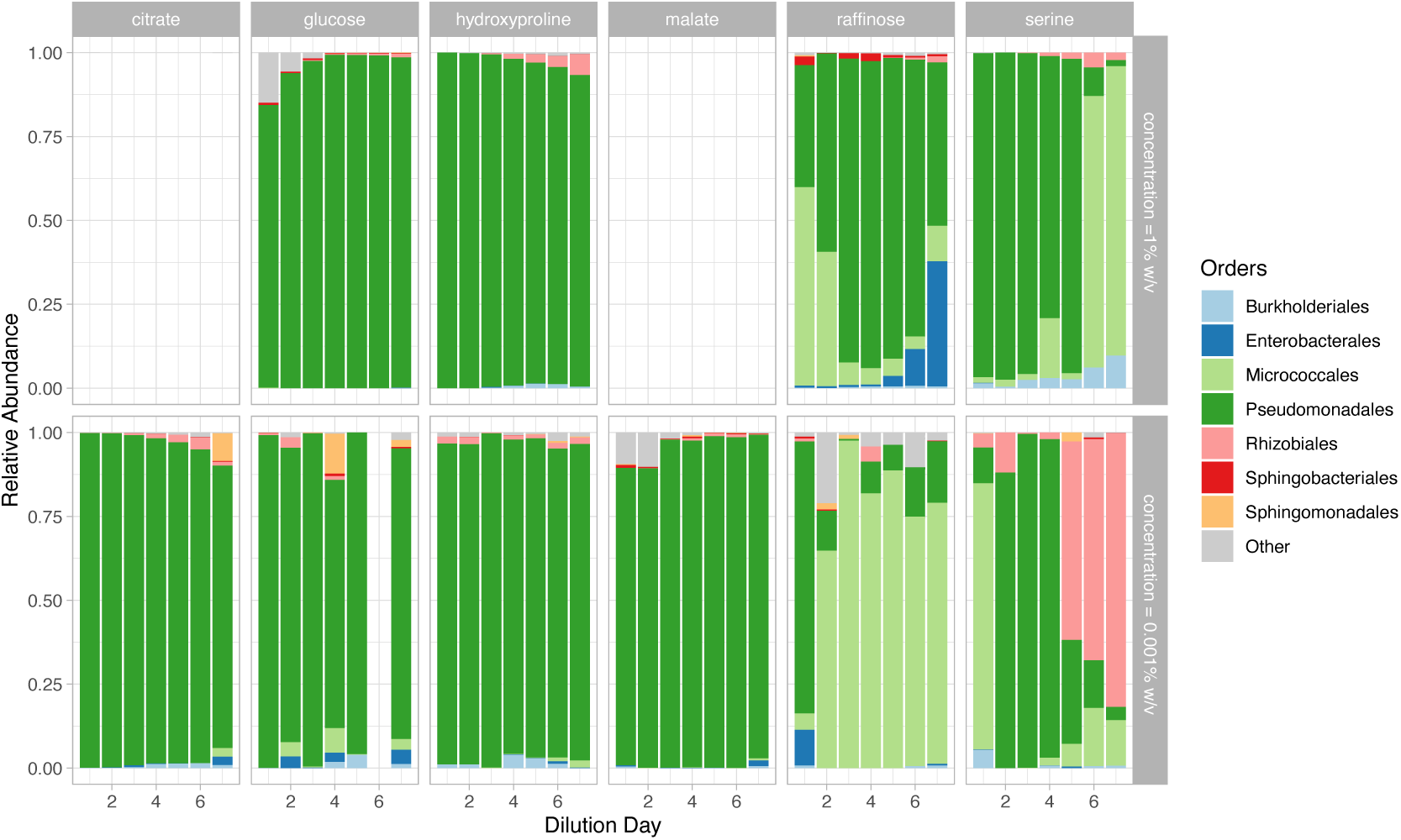
The majority of communities reached equilibrium before the end of the experiment. Each panel shows the temporal trajectories of the composition of one community at the order level. The 8 most prevalent orders are included. We sequenced twelve time series in total, including six carbon sources per two concentrations.

**Figure S2:**
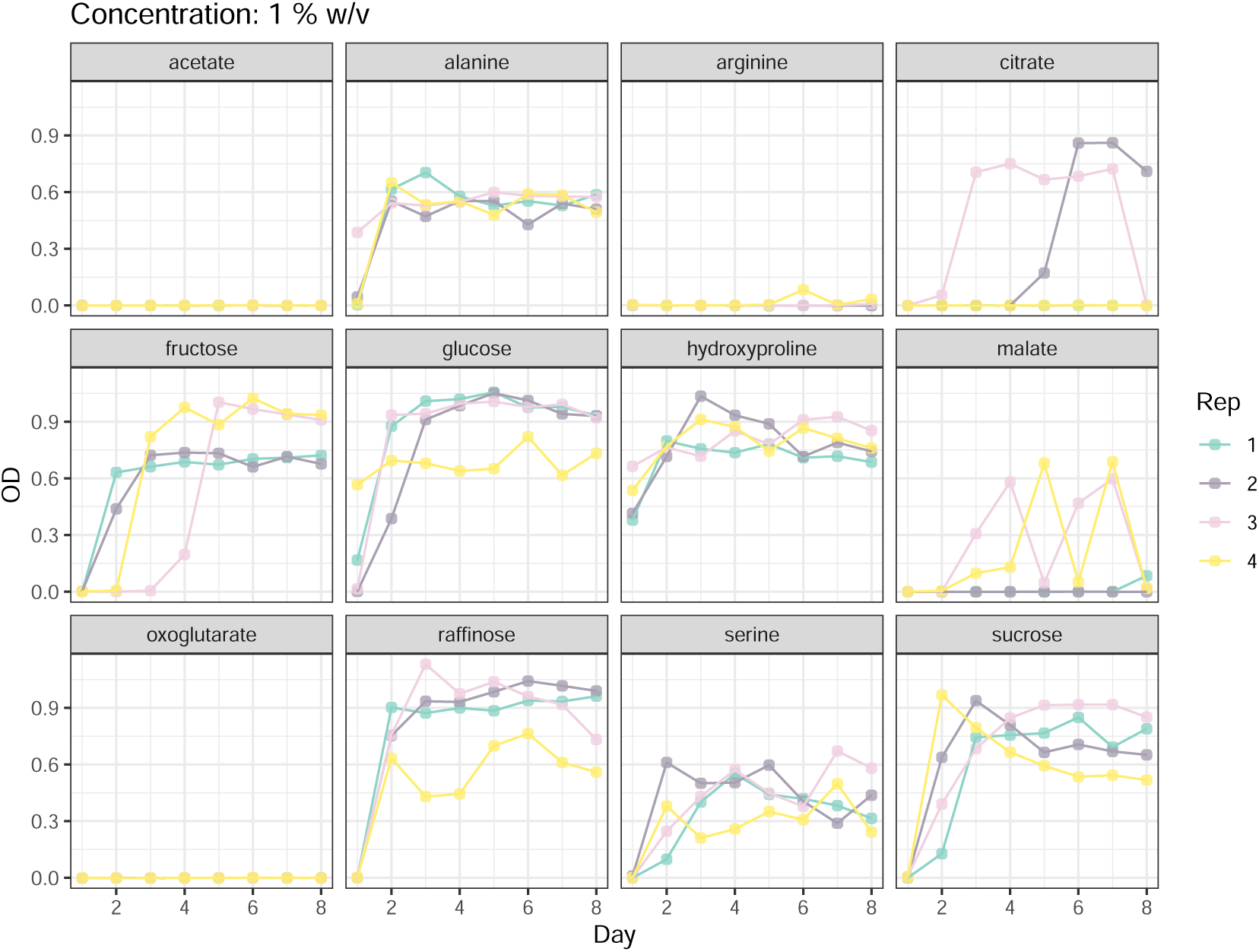
Biomass trajectories of communities grown at a carbon concentration of 1% w/v. All carbon sources except organic acids (citrate, acetate, malate, oxoglutarate) and the amino acid arginine sustained dense communities. Replicated communities grown in sugars (glucose, fructose, sucrose, raffinose) displayed substantial variation in biomass, possibly indicative of the presence of alternative states. Communities grown on organic acids likely collapsed due to the low pH of the growth medium. Biomass was measured as OD600, optical density at 600 nm, in 80 *µ*l of culture.

**Figure S3:**
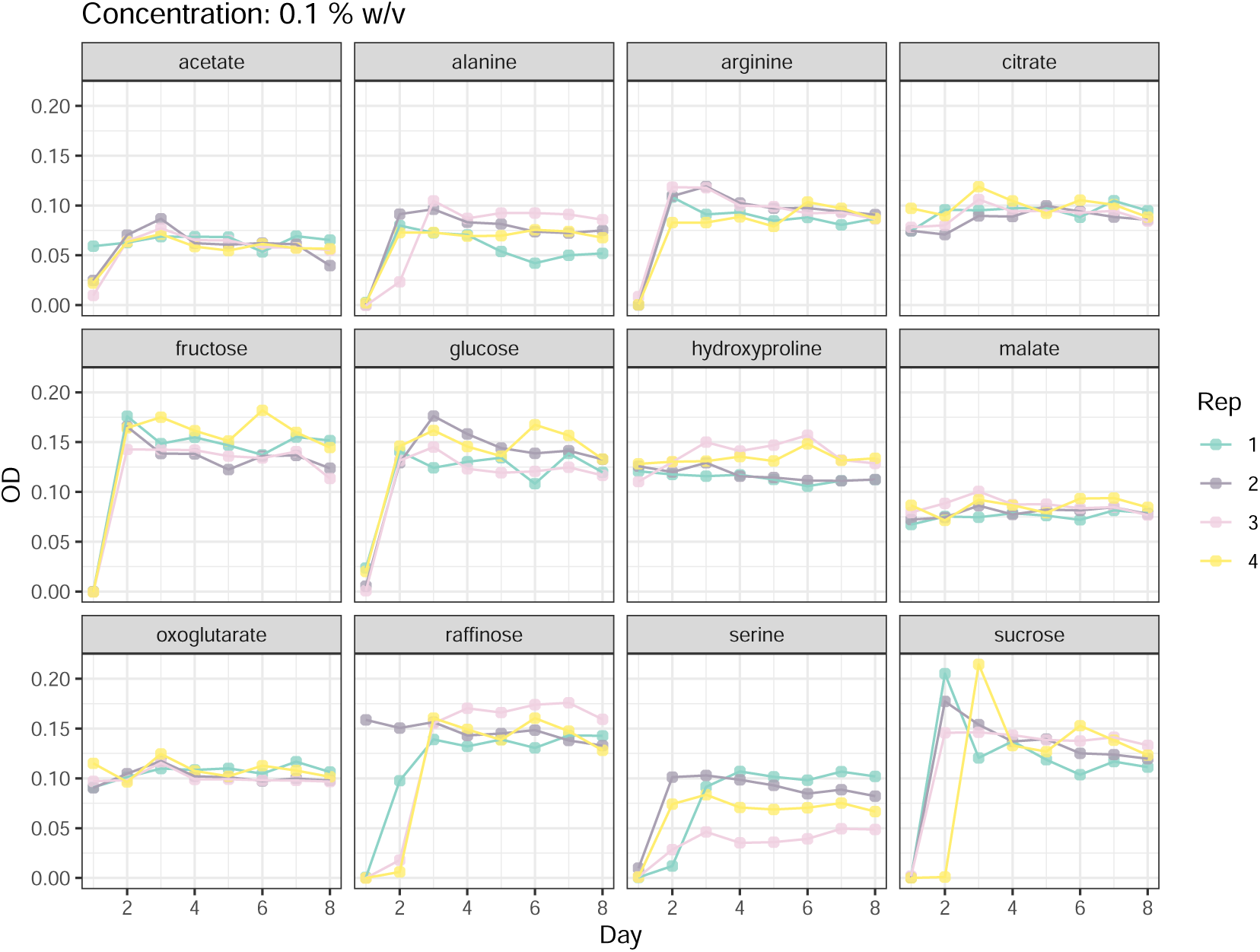
Biomass trajectories of communities grown at a carbon concentration of 0.1% w/v. Biomass was measured as OD600, optical density at 600 nm, in 80 *µ*l of culture.

**Figure S4:**
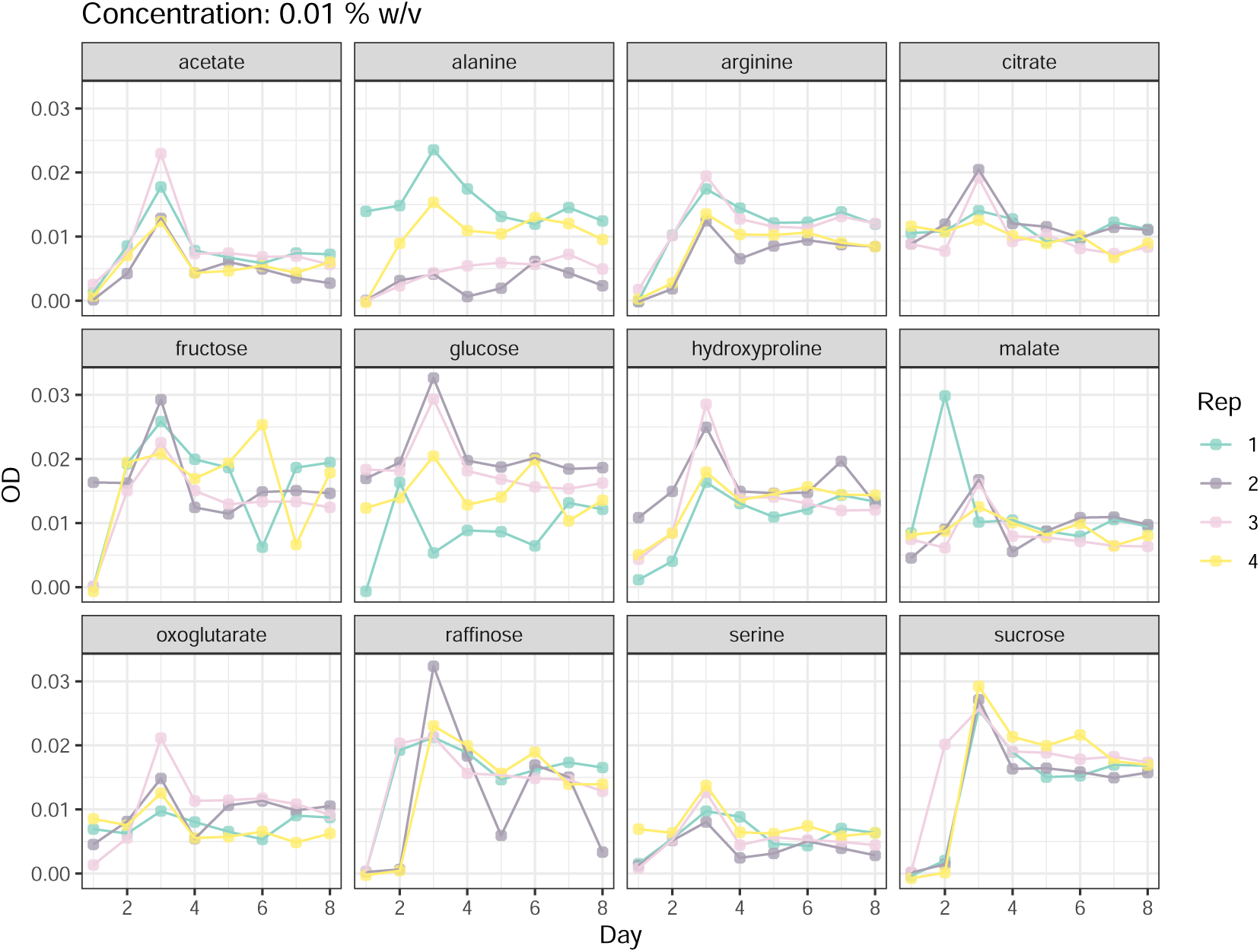
Biomass trajectories of communities grown at a carbon concentration of 0.01% w/v. Biomass was measured as OD600, optical density at 600 nm, in 80 *µ*l of culture.

**Figure S5:**
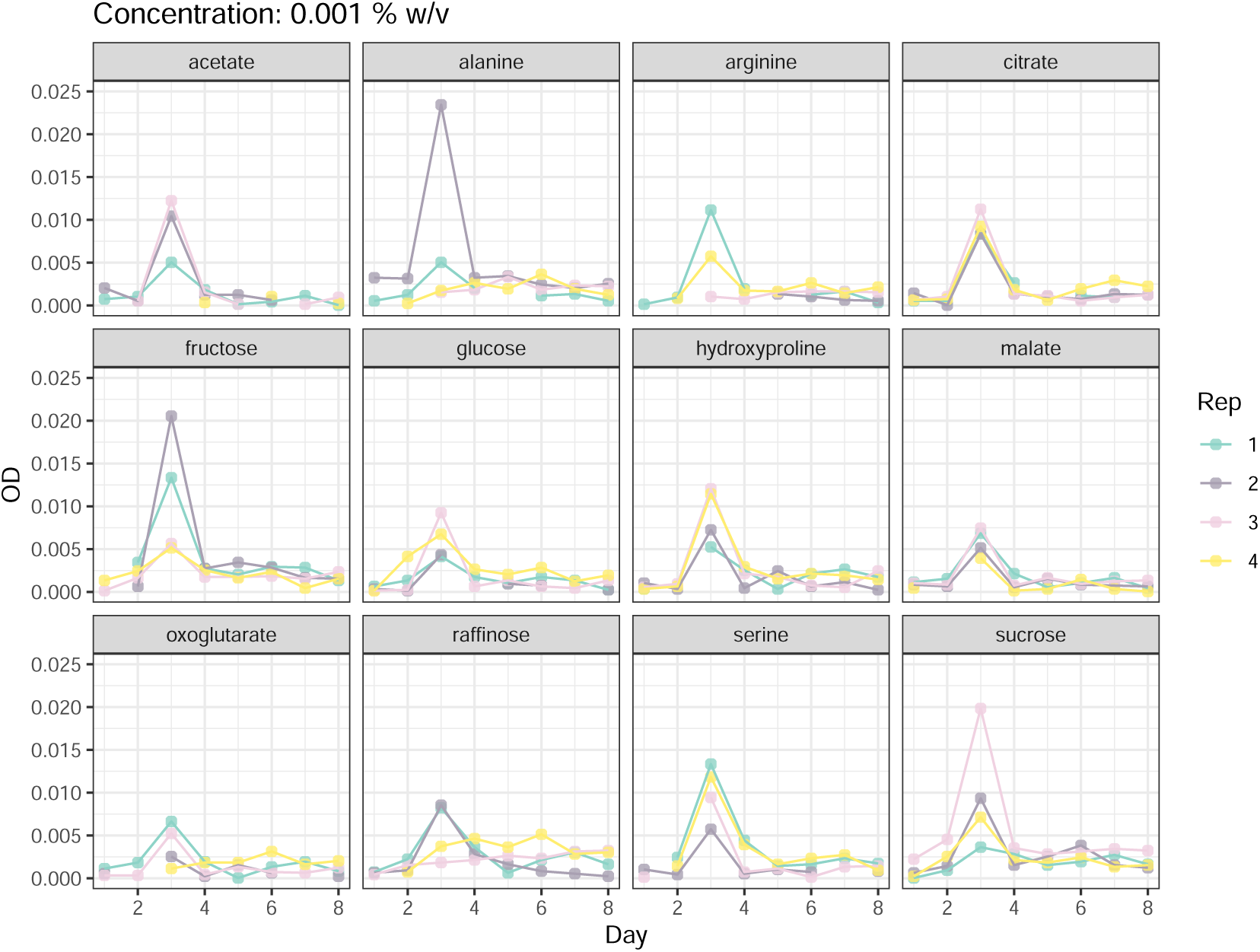
Biomass trajectories of communities grown at a carbon concentration of 0.001% w/v. Biomass was measured as OD600, optical density at 600 nm, in 80 *µ*l of culture.

**Figure S6:**
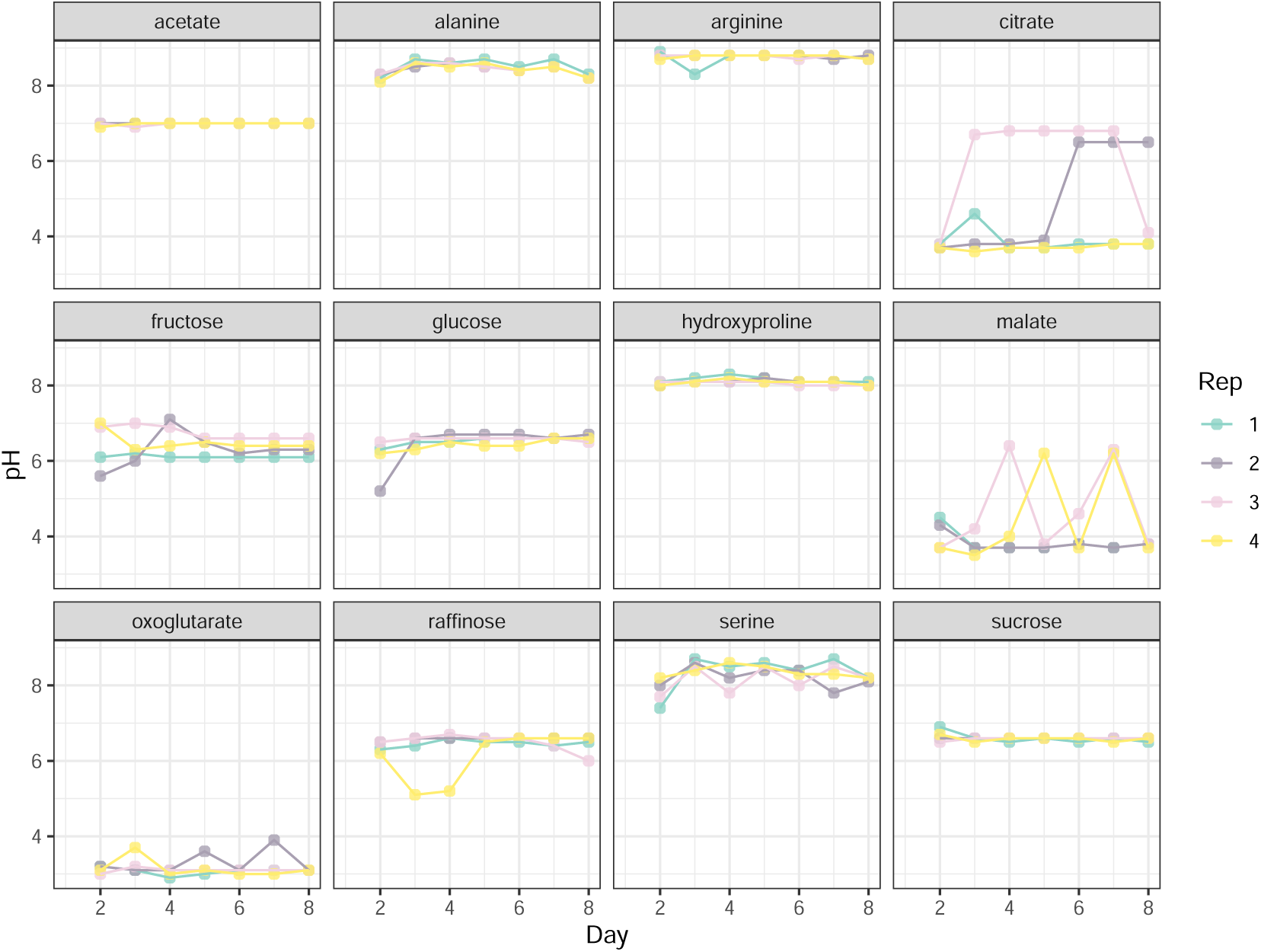
pH trajectories of communities grown at a carbon concentration of 1% w/v. The majority of collapsed communities displayed low pH during the entire experiment.

**Figure S7:**
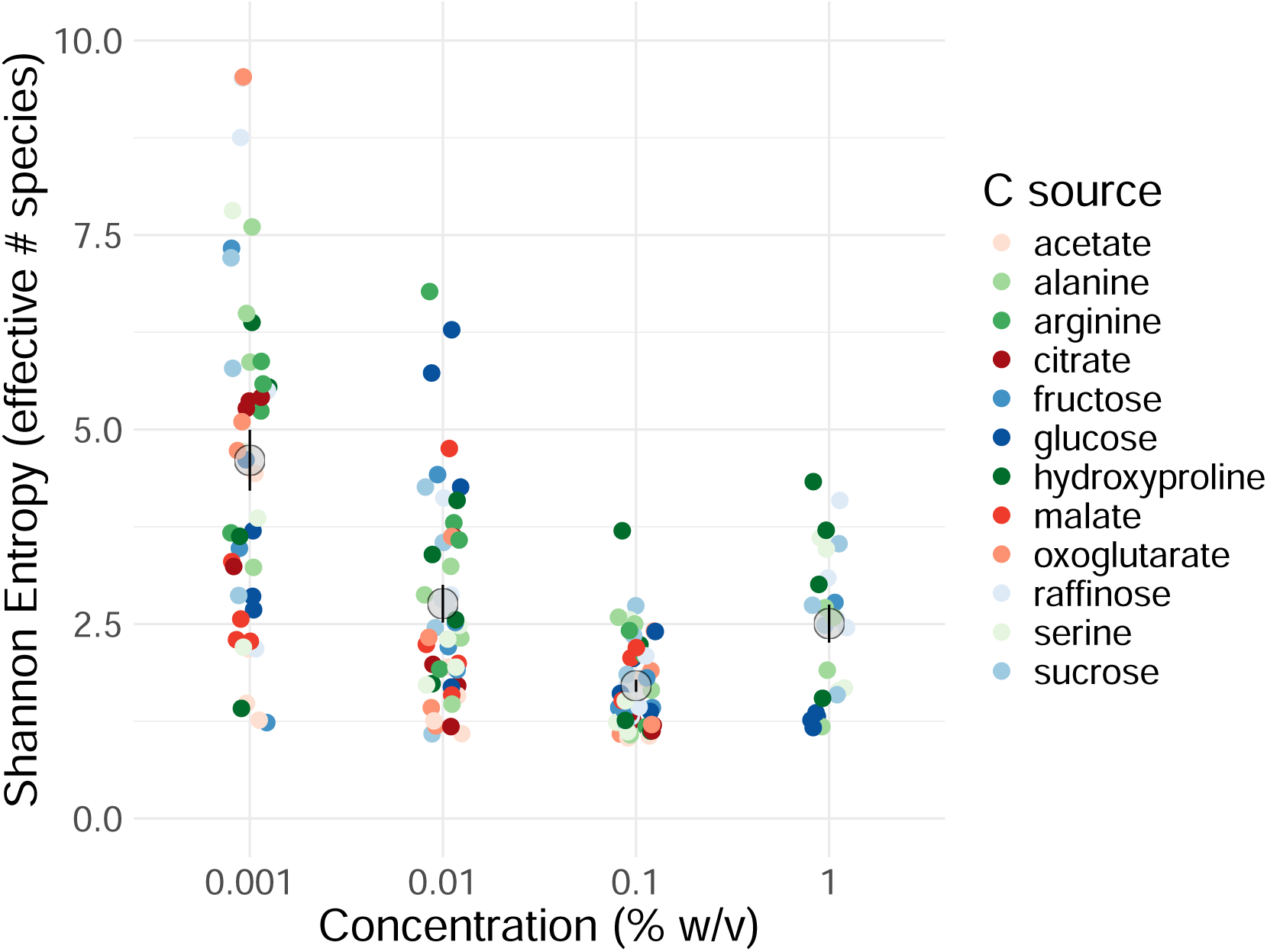
Diversity, measured as Shannon Entropy slightly decreases with carbon source concentration. The effective number of species is plotted as a function of resource concentration. Shannon entropy is calculated for all replicates and averaged for each concentration.

**Figure S8:**
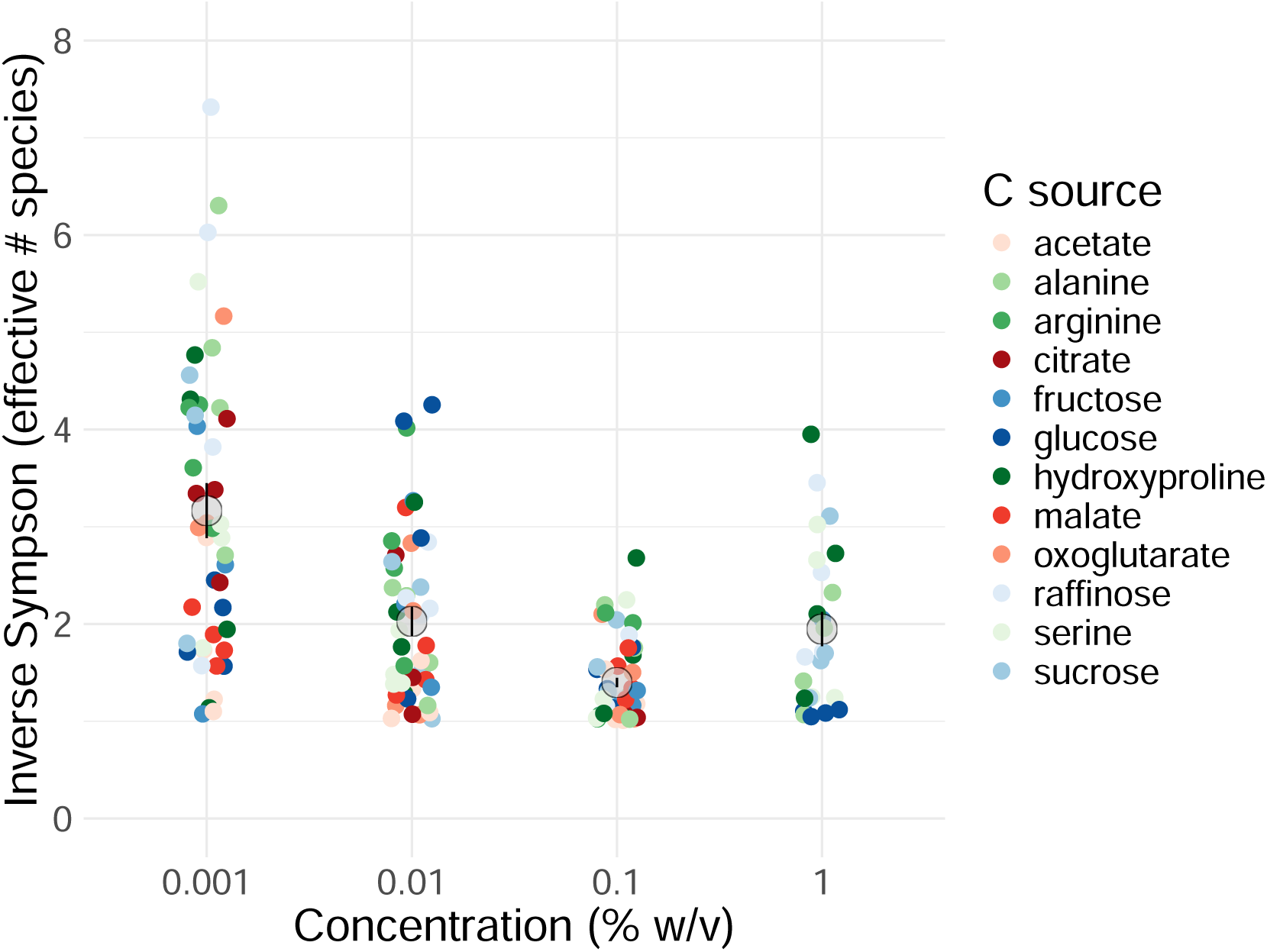
Diversity, measured as Inverse Simpson index slightly decreases with carbon source concentration. The effective number of species is plotted as a function of resource concentration. Inverse Simpson is calculated for all replicates and averaged for each concentration.

**Figure S9:**
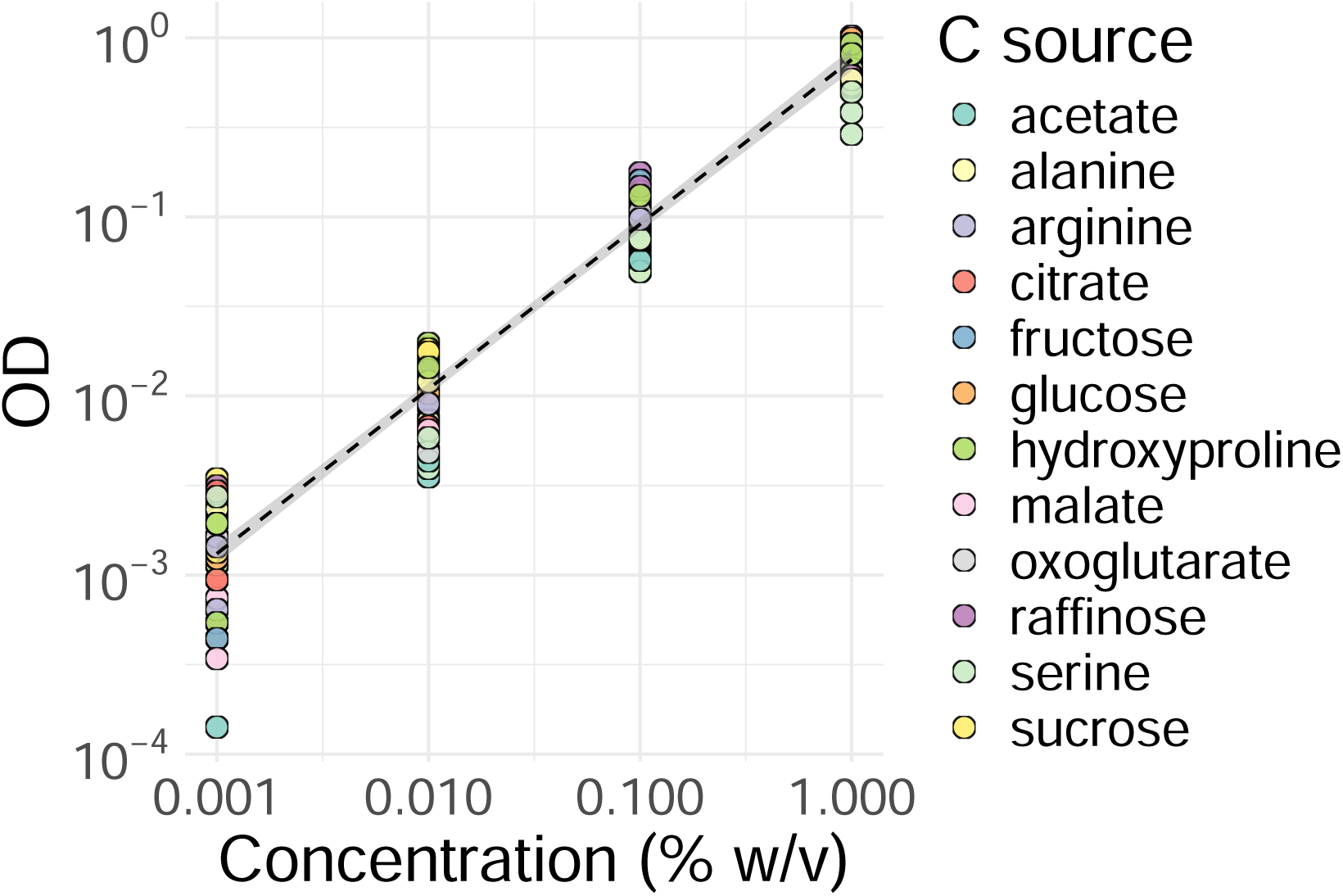
Biomass of communities increases with the concentration of the supplied carbon source. Biomass was measured as optical density (OD600) at the end of each dilution day. Here we plotted OD600 at day 7.

**Figure S10:**
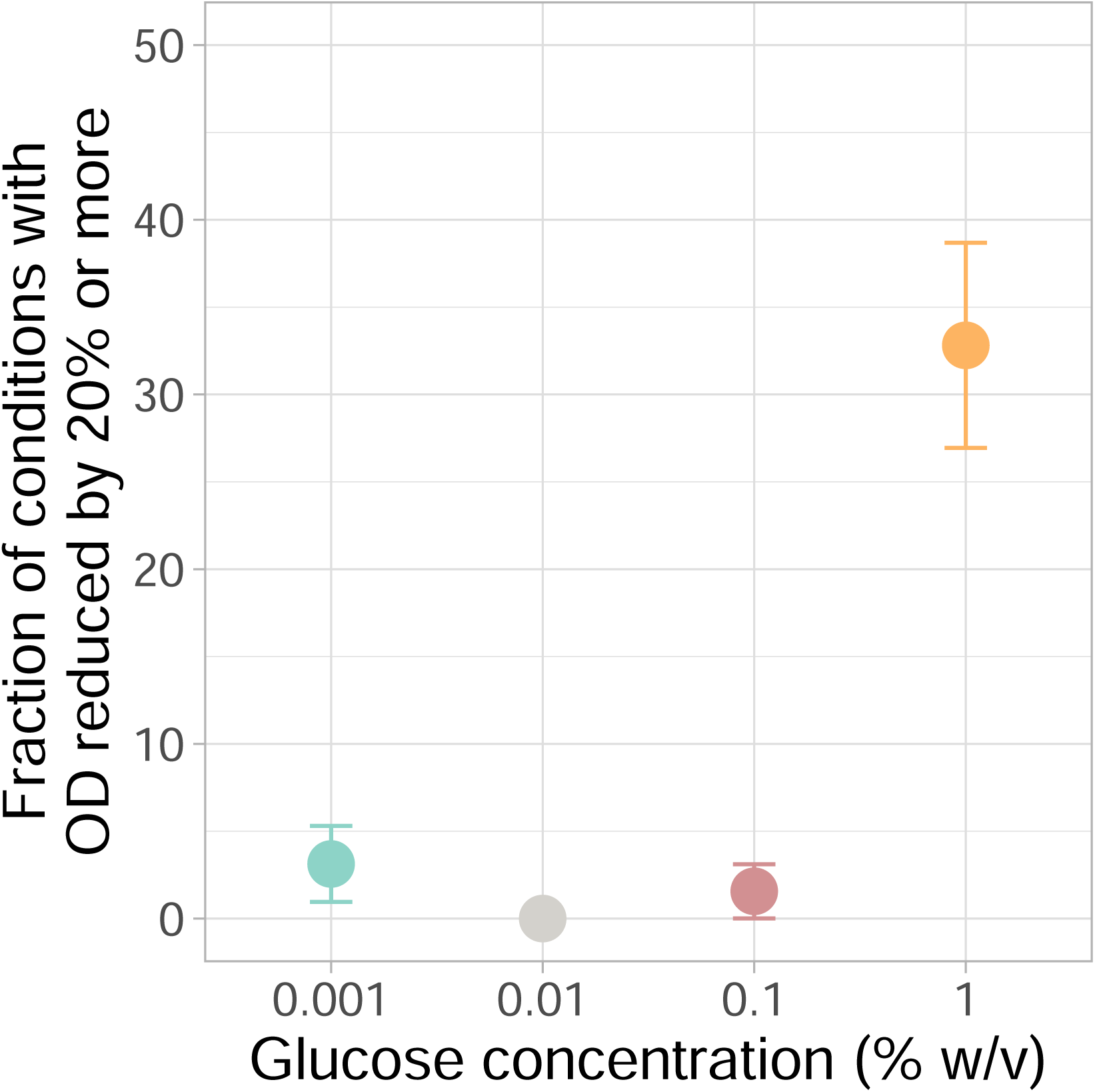
More than 30% of species experienced a reduction of 20% in final optical density in spent media obtained at the highest concentration of carbon. Fraction of conditions, i.e. individual species cultures in glucose supplemented with spent media from 8 species, showing a reduction in final optical density of 20% as a function of the original concentration of glucose. The binomial standard error is displayed (N = 64 for each glucose concentration)

**Figure S11:**
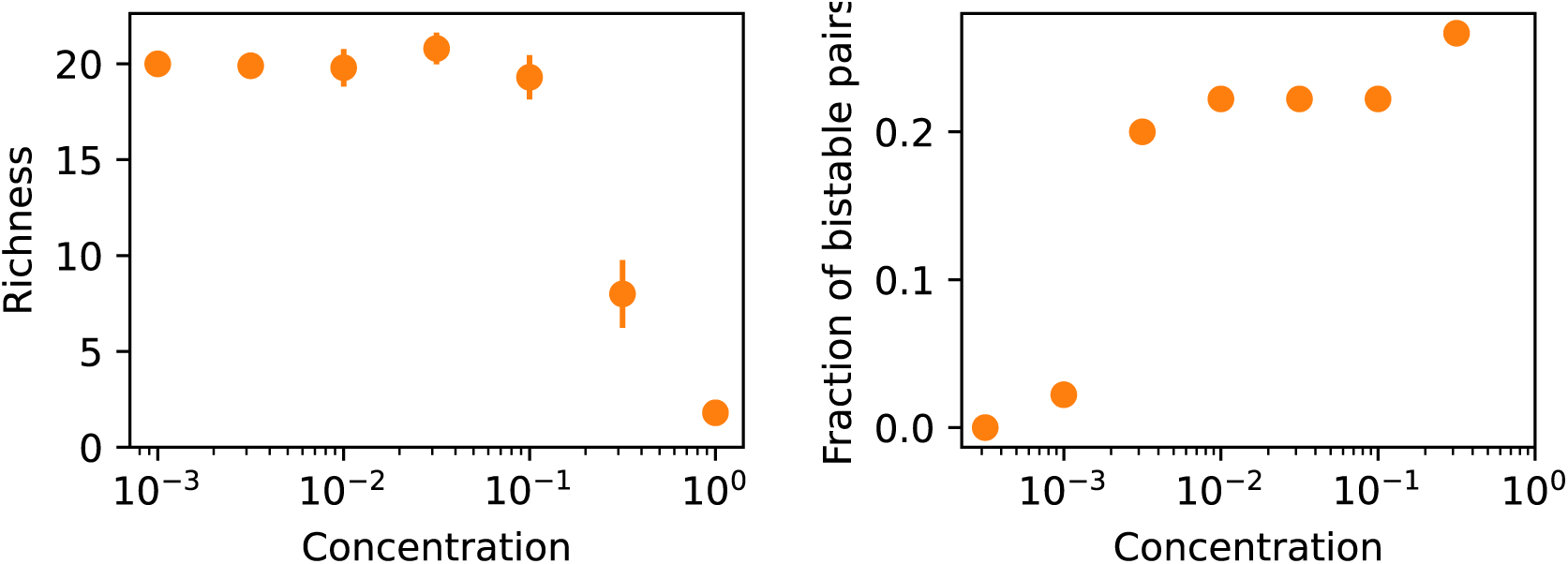
The patterns due to toxicity do not change with more complicated ways of setting the toxicity threshold. *Q*. In the main text, for simplicity, we assume a fixed *Q* for all species and resources. If the flux *J_iα_* is close to *Q*, the yield for species *i* on resource *α* will decrease. Here, we make the threshold depend on species and resources, sampling a *Q_iα_* from a uniform distribution U[0.75*Q,* 1.25*Q*], where *Q* is the mean value. We next simulated the communities with all other parameters the same as Fig. 5. We found that richness decreases at a similar supplied concentration (Fig. 5F), and the fraction of bistable pairs follows a similar relationship with concentration as Fig. 5G.

**Figure S12:**
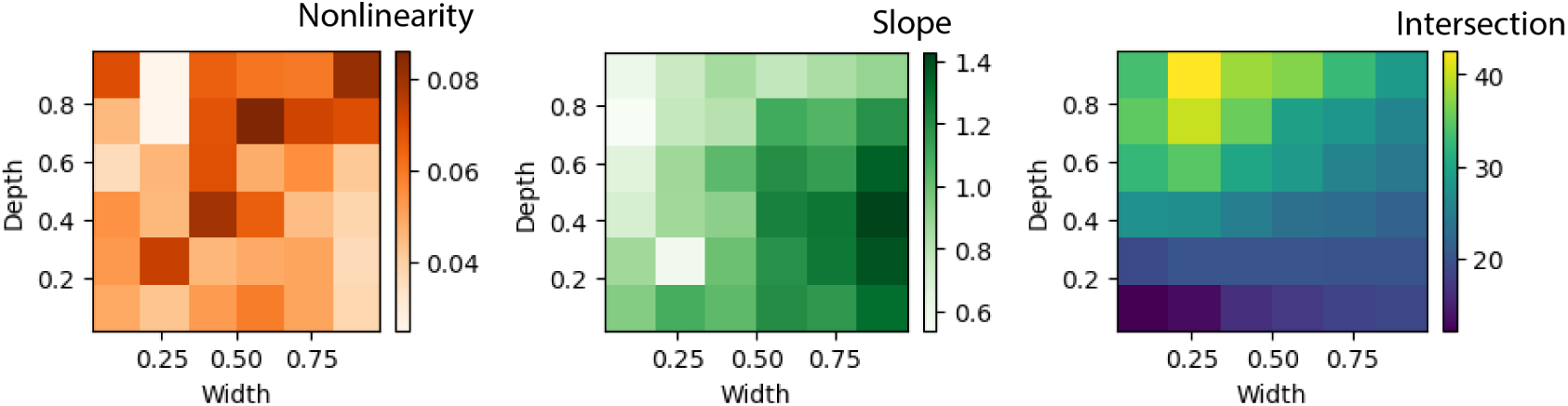
Adding toxicity does not affect richness patterns for wide and shallow networks at low supplied concentrations. After adding toxicity to the model as Fig. 5, we fix the concentration as Fig. 3 and study the relationship between richness and number of supplied resources again. For wide and shallow networks, we observe small nonlinearity, a linear slope close to 1, and an intersection (richness with one supplied resource) around 20, which are similar to the case without toxicity. The intuition is that for each resource, the concentration is small, which has not yet reached the threshold of being toxic. However, under the same supplied concentration, narrow networks are more susceptible to toxicity, being different from the simulation without toxicity. An explanation is that narrow networks will produce metabolites with higher concentration, which can be more toxic given the same supply flux.

**Figure S13:**
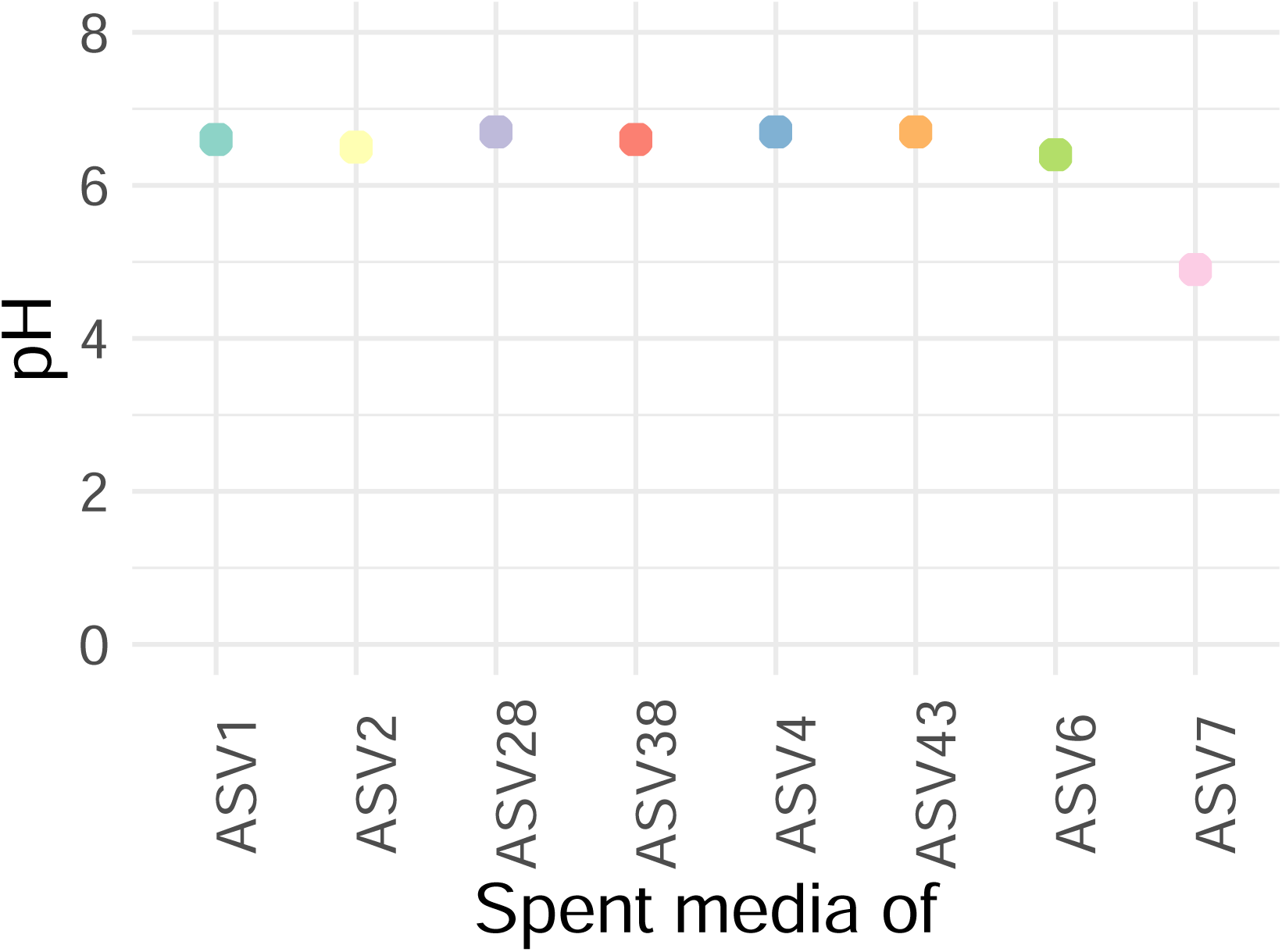
pH trajectories of the spent media of the 8 isolates grown at glucose 1% w/v. The majority of collapsed communities displayed low pH during the entire experiment.

**Figure S14:**
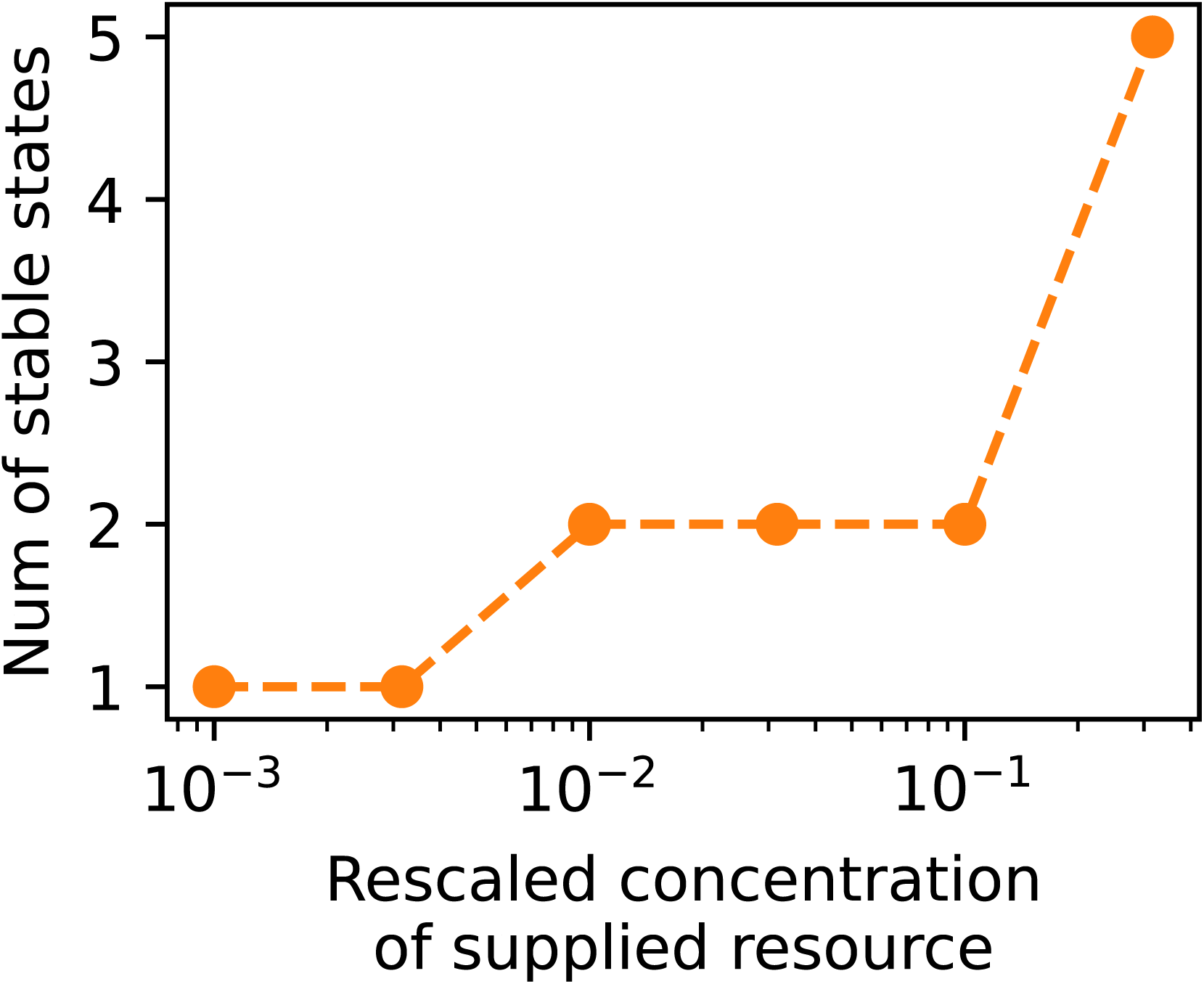
Model confirms that the number of stable states in a community increases with resource concentration. Simulated communities show a loss of global stability when the supplied resource concentration exceeds a threshold. The number of stable states is obtained by PCA analysis of the final abundance vectors obtained from different initial conditions. The threshold where the community begins to have alternative stable states agrees with that of where bistable pairs emerge.

**Figure S15:**
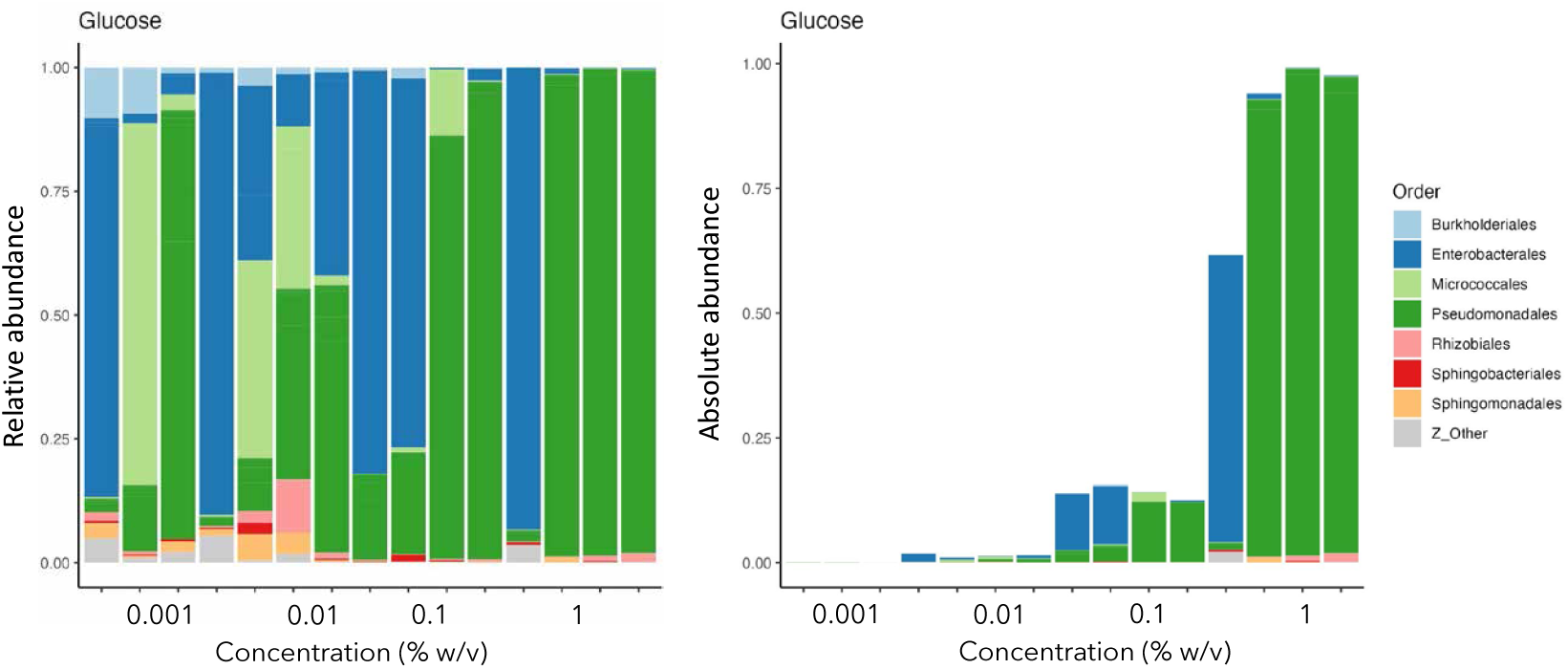
Two distinct biomass states, each dominated by a different taxon, are identified in communities grown in glucose supplemented at 1% w/v. Composition of communities grown in M9 supplemented with glucose (all replicates for each concentration are shown). Left panel, relative abundance. Right panel, absolute abundance obtained by multiplying relative abundance by the final OD of the community at day 7.

## Notes

### Competing Interest Statement

The authors have declared no competing interest.

